# Distinct fibrotic, epithelial and immune transcriptomic programs in phenotypes of chronic lung allograft dysfunction

**DOI:** 10.64898/2026.05.24.727536

**Authors:** Tsukasa Ishiwata, Gregory Berra, Jonathan Allen, Ankita Burman, Gavin Wilson, Zoeen Carter, Tatsuaki Watanabe, Melinda Solomon, Shaf Keshavjee, Jonathan Yeung, Stephen Juvet, Tereza Martinu

## Abstract

**Background:** Chronic lung allograft dysfunction (CLAD) is the major cause of late mortality after lung transplantation and includes two principal phenotypes, bronchiolitis obliterans syndrome (BOS) and restrictive allograft syndrome (RAS). RAS and other phenotypes with RAS-like opacities (RLO) on chest imaging have a poorer prognosis. Despite clear clinical and pathological differences, molecular distinctions between phenotypes remain poorly defined. We aimed to explore gene transcriptional profiles across CLAD phenotypes and relevant controls.

**Methods:** We performed bulk RNA sequencing on explanted lung tissue from 45 lung transplant recipients with end-stage CLAD (20 with RLO and 25 without RLO). Samples from twenty-seven control donor and lobectomy lungs and sixteen idiopathic pulmonary fibrosis (IPF) lungs served as comparators. Non-negative matrix factorization (NMF) was used to identify latent transcriptomic signatures, which were correlated with clinical, radiologic, and histopathologic features.

**Results:** NMF identified seven distinct gene signatures that segregated CLAD phenotypes. RLO-CLAD lungs were enriched for extracellular matrix remodeling and B-cell/plasma cell–associated signatures, overlapping partly with IPF, whereas non-RLO-CLAD showed relative enrichment of epithelial injury and surfactant-response pathways. Signatures related to epithelial homeostasis and ciliary/microtubule function were progressively reduced from control lungs to non-RLO-CLAD and were most suppressed in RLO-CLAD.

**Conclusions:** RLO-CLAD and non-RLO-CLAD, aligning with RAS and BOS phenotypes, show distinct transcriptomic signatures. RLO-CLAD is characterized by profibrotic and humoral immune signatures with profound epithelial dysfunction, whereas non-RLO-CLAD shows relative enrichment of epithelial injury responses. These data provide molecular stratification of CLAD and support the development of phenotype-specific biomarkers and targeted therapies.

## Background

Lung transplantation (LTx) is considered the ultimate treatment for carefully selected patients with end-stage lung disease, for whom no other effective therapeutic options remain. However, the long-term success after LTx is significantly hampered by chronic lung allograft dysfunction (CLAD), a progressive and irreversible decline in lung function that is the leading cause of morbidity and mortality beyond the first year post-LTx.^1^ CLAD is clinically classified into multiple types, including two principal phenotypes: bronchiolitis obliterans syndrome (BOS), characterized by obstructive lung physiology due to small-airway obliteration, and restrictive allograft syndrome (RAS), defined by restrictive pulmonary physiology and diffuse parenchymal fibrosis with RAS-like opacities (RLO) on chest imaging.^2–5^ Patients may also present with a mixed phenotype (with obstruction, restriction and RLO) or with undefined or unclassified phenotypes (with or without RLO).^2,6^ CLAD patients can also transition between phenotypes over time.^7^ Notably, RAS and mixed phenotypes constitute about 14-32% of CLAD cases and carry a dramatically worse prognosis. Median survival after RAS onset is on the order of 18 months versus nearly 4 years for BOS.^2,5^ Importantly, undefined or unclassified CLAD phenotypes with RLO show survival outcomes similar to those of RAS or mixed phenotypes.^6^ This indicates that survival risk can be stratified based on the presence of RLO on chest imaging, and that the RLO-CLAD phenotype is associated with poor survival outcomes. Pathologically, BOS is defined by airway-centered obliterative bronchiolitis, whereas RLO-CLAD phenotypes demonstrate widespread involvement of alveoli and pleura with patterns of pleuroparenchymal fibroelastosis, acute fibrinous and organizing pneumonia, diffuse alveolar damage, and other fibrotic lesions.^3–5,8^

Despite these well-defined clinical and pathological distinctions, the precise molecular mechanisms that differentiate RLO-CLAD from non-RLO-CLAD remain poorly understood. Current knowledge gaps include a comprehensive understanding of the specific cellular and molecular pathways that drive the divergent fibrotic and inflammatory processes in each phenotype. Addressing these gaps is crucial for the development of phenotype-specific, or perhaps endotype-specific, diagnostic tools and targeted therapeutic interventions, moving beyond standard-of-care immunosuppression. Previous research has highlighted the involvement of alloimmune responses, external stimuli (e.g., infection, aspiration), and genetic variants in CLAD pathogenesis,^9–11^ but a clear molecular stratification of RLO-CLAD and non-RLO-CLAD has been elusive. This study aims to address this critical knowledge gap by performing bulk RNA sequencing on explanted lung tissue from CLAD patients. By applying non-negative matrix factorization (NMF), a computational method designed to identify underlying biological processes from complex gene expression data, this investigation seeks to uncover distinct transcriptomic signatures that differentiate the more fibrotic and severe RLO-CLAD from non-RLO-CLAD phenotypes.

## Methods

### Patient Cohort and Sample Collection

This study was conducted in accordance with ethical guidelines and received institutional approval under protocol numbers #15-9531 and #21-5551. Prior to any sample collection, informed written consent was obtained from all patients or their legal representatives.

Explanted lung tissue specimens were prospectively collected at Toronto General Hospital from a cohort of 45 LTx recipients diagnosed with end-stage CLAD who underwent redo LTx or autopsy between May 2015 and September 2020. All included patients met the ISHLT consensus criteria for CLAD.^1^ For CLAD samples, the right upper lobe was selected when available; otherwise, the left upper lobe was used. Both central and peripheral lung tissue was collected to ensure comprehensive representation of the heterogeneous disease pathology, as previously described.^12^

Additional lung tissue samples were included as controls: Donor lung tissue (n=17) was collected at the end of cold ischemia from discarded portions of downsized donor lungs (15 from donation after brain death, 2 from donation after circulatory death). Donor lungs that were declined for transplantation were excluded. Additional control samples were obtained from patients undergoing lobectomy for malignancy (n=10), from areas remote from the suspected cancer. Strict inclusion criteria were applied, ensuring that these lobectomy patients had normal pulmonary function, and had no pathological evidence of cancer or significant lung disease in the sampled tissue.

Given the fibrotic nature of RAS, idiopathic pulmonary fibrosis (IPF) lungs (n=16) were included as a relevant positive comparator. These samples were collected at the time of LTx from ‘untreated’ IPF patients, defined as those not receiving immunosuppressive therapy (prednisone ≤10 mg/day allowed) and free from anti-fibrotic agents (nintedanib or pirfenidone) for at least 6 months prior to LTx. To ensure a well-defined IPF phenotype, patients with alternative diagnoses or a recent acute exacerbation within the past month were excluded.

### CLAD Phenotyping

CLAD patients’ phenotypes were adjudicated at time of lung explant as BOS, RAS, mixed, or undefined based primarily on the ISHLT consensus definitions,^1^ while the remaining cases were labelled as “unclassified” CLAD.^6^ RLO on chest imaging at CLAD onset were carefully identified—defined as opacities and/or increasing pleural thickening consistent with pulmonary and/or pleural fibrosis and likely to cause restrictive physiology—to capture the radiologic findings associated with fibrotic remodeling and lower survival. Following this CLAD phenotyping, samples were grouped a priori into four distinct categories for downstream RNA sequencing analysis: ‘RLO-CLAD’, ‘non-RLO-CLAD’, ‘IPF’, and ‘control’.

### RNA-sequencing Library Preparation and Processing

To ensure the integrity and quality of RNA, all tissue samples were processed within 6 hours of explant or resection. For CLAD lungs, samples approximately 2 cm² in size were immediately immersed in RNAlater (QIAGEN) at 4°C for 24 hours, after which the RNAlater was decanted, and samples were stored at -80°C until further processing. For IPF and control lungs, samples were snap-frozen in liquid nitrogen and then stored at -80°C until further processing. Total RNA was extracted from the prepared tissue samples using TRIzol (Thermo Fisher), and its quality was assessed to confirm suitability for high-throughput sequencing. RNA with appropriate RNA integrity number scores (average >7) was used to prepare libraries. RNA-sequencing libraries were generated utilizing the NEBNext Ultra II Directional RNA Library Prep Kit with poly-A selection, specifically targeting messenger RNA (mRNA) to capture protein-coding gene expression profiles. Sequencing was performed on an Illumina NovaSeq platform (S2 flow cell, 100-cycle), aiming to achieve approximately 50 million 1x100 bp single-end reads per sample.

### BulkRNAseq Data Analysis

For analyses where only one sample per patient was desired, data from peripheral lung regions of CLAD samples were preferentially used when both central and peripheral samples were available.

#### Gene expression quantification

Quantifying gene expression using Salmon followed by tximport to calculate gene level transcripts per million (TPM). Top 5% protein-coding gene TPMs with the highest variance were used in NMF.

#### Principal Component Analysis

To evaluate the global transcriptomic relationships among the different lung phenotypes, we performed a Principal Component Analysis (PCA) using the top 5000 most variable genes across all samples.

#### Non-negative Matrix Factorization (NMF) Analysis

NMF was employed as a computational method to identify coherent gene expression patterns, or signatures, within the normalized expression matrix. The NMF algorithm decomposes the gene-by-sample matrix V into two non-negative matrices: W (metagenes, representing the expression profiles of underlying biological processes) and H (sample weights, indicating the contribution of each signature to a given sample’s overall expression profile), such that V ≈ WH. The number of factors (k) ranging from 2 to 20 were evaluated, and k = 7 was selected based on the reconstruction error analysis. Signature score indicates how strongly a given sample expresses a particular NMF-derived gene signature (i.e., a higher signature score for a sample means that the sample’s transcriptomic profile closely aligns with the gene pattern represented by that signature). The top 50 genes ranked in each identified signature based on the weights in the W matrix were subsequently interpreted through Gene Ontology (GO) and pathway analysis to infer the associated biological functions. Signatures with significantly higher expression in RLO-CLAD compared to non-RLO-CLAD were defined as RAS-enriched, and vice versa, based on a Wilcoxon test with a false discovery rate adjusted p-value <0.05.

### SCGB3A2 Protein in Bronchoalveolar Lavage

SCGB3A2 protein levels were measured in bronchoalveolar lavage (BAL) fluid from an independent cohort.^13^ BAL samples were obtained from 76 LTx recipients: 53 with CLAD (26 RLO-CLAD and 27 non-RLO-CLAD) and 23 stable controls without CLAD. CLAD samples were collected within ±3 months of clinical diagnosis. SCGB3A2 was quantified by ELISA (CSB-EL020819HU-5, Cusabio, TX, USA). Total protein was measured using bicinchoninic acid total protein assay. SCGB3A2 concentrations were normalized to total protein.

### Ciliated Cell Composition

In a subset of the bulk RNA sequencing cohort, samples from 29 lungs (8 RLO-CLAD LTx, 11 non-RLO-CLAD LTx, and 10 donor controls) were analyzed, and airway epithelial composition was assessed by immunofluorescence. Formalin-fixed paraffin-embedded sections were stained for pan-cytokeratin (pan-CK) (1:100 anti-pan-CK [ab9377, Abcam, Cambridge, UK] and 1:1000 Donkey anti-rabbit IgG plus 488 [A32790, Thermo Fisher Scientific, MA, USA]) and the ciliated cell marker FOXJ1 (1:100 anti-FOXJ1 [14-9965-82, Thermo Fisher Scientific] and 1:1000 Goat anti-mouse IgG plus 555 [A21127, Thermo Fisher Scientific]). Pan-CK^+^ epithelial cells and FOXJ1^+^ pan-CK^+^ ciliated cells were quantified using HALO image analysis software (v3.0311, Indica Labs, NM, USA). The percentage of ciliated cells (FOXJ1^+^ pan-CK^+^ cells among all pan-CK^+^ epithelial cells) was compared between RLO-CLAD and non-RLO-CLAD groups.

### Clinical Correlations with NMF-Derived Gene Signatures

Associations between gene signatures and clinical variables, including CLAD phenotypes, radiologic and histopathologic findings, and clinical donor/recipient characteristics, including age, sex, smoking history, cytomegalovirus (CMV) mismatch risk, acute rejection score,^14^ infection score,^15^ bacterial/fungal colonization or infection at time of explant, and donor-specific antibody (DSA) history, were assessed using Spearman correlation.

## Results

### Patient Characteristics and Clinical Data

The clinical and demographic characteristics of the study cohort are summarized in Table 1. A total of 88 subjects were included, comprising patients with CLAD (n=45), IPF (n=16), and control lungs (n=27) (17 donor lungs, 10 lobectomy lungs). Among the patients with CLAD, 20 (44.5%) were classified as RLO-CLAD and included RAS, mixed, and undefined phenotypes (which all had RLO). All other CLAD cases were non-RLO-CLAD (n=25, 55.6%) and included BOS and unclassified cases (all of which had no RLO).

**Table 1.**
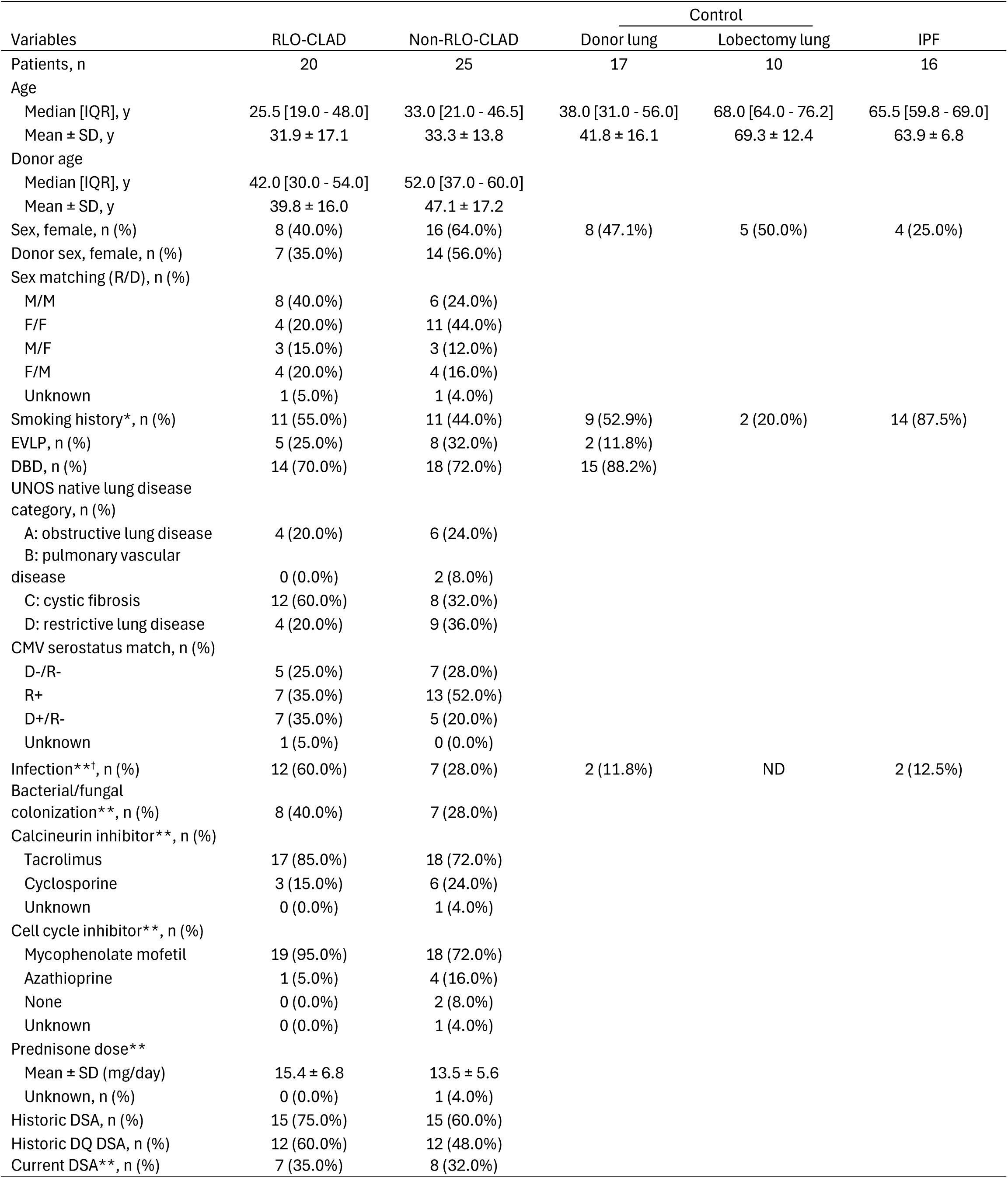
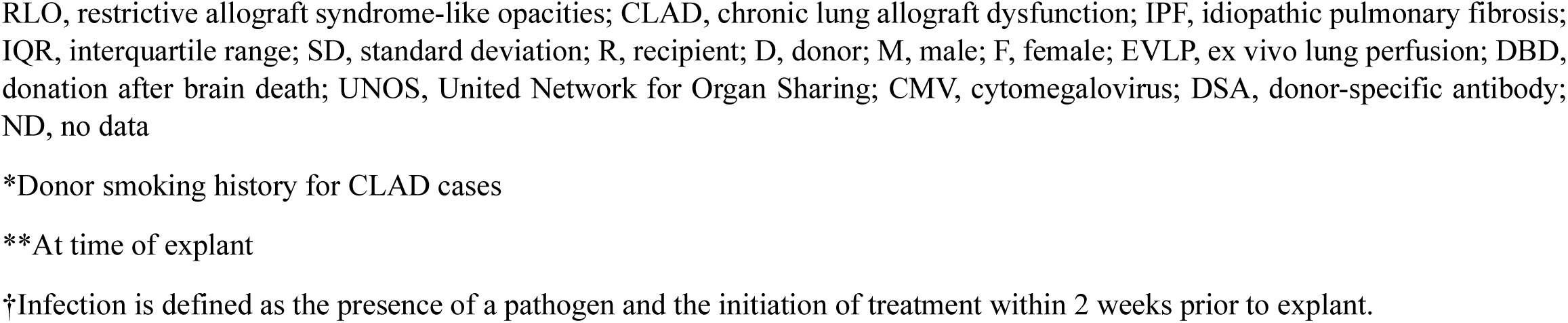
Clinical characteristics of the study subjects.

| Variables | RLO-CLAD | Non-RLO-CLAD | Control |  | IPF |
| --- | --- | --- | --- | --- | --- |
|  |  |  | Donor lung | Lobectomy lung |  |
| Patients, n | 20 | 25 | 17 | 10 | 16 |
| Age |  |  |  |  |  |
| Median [IQR], y | 25.5 [19.0 - 48.0] | 33.0 [21.0 - 46.5] | 38.0 [31.0 - 56.0] | 68.0 [64.0 - 76.2] | 65.5 [59.8 - 69.0] |
| Mean ± SD, y | 31.9 ± 17.1 | 33.3 ± 13.8 | 41.8 ± 16.1 | 69.3 ± 12.4 | 63.9 ± 6.8 |
| Donor age |  |  |  |  |  |
| Median [IQR], y | 42.0 [30.0 - 54.0] | 52.0 [37.0 - 60.0] |  |  |  |
| Mean ± SD, y | 39.8 ± 16.0 | 47.1 ± 17.2 | | | |
| Sex, female, n (%) | 8 (40.0%) | 16 (64.0%) | 8 (47.1%) | 5 (50.0%) | 4 (25.0%) |
| Donor sex, female, n (%) | 7 (35.0%) | 14 (56.0%) |  |  |  |
| Sex matching (R/D), n (%) |  |  |  |  |  |
| M/M | 8 (40.0%) | 6 (24.0%) |  |  |  |
| F/F | 4 (20.0%) | 11 (44.0%) |  |  |  |
| M/F | 3 (15.0%) | 3 (12.0%) |  |  |  |
| F/M | 4 (20.0%) | 4 (16.0%) |  |  |  |
| Unknown | 1 (5.0%) | 1 (4.0%) |  |  |  |
| Smoking history*, n (%) | 11 (55.0%) | 11 (44.0%) | 9 (52.9%) | 2 (20.0%) | 14 (87.5%) |
| EVLP, n (%) | 5 (25.0%) | 8 (32.0%) | 2 (11.8%) |  |  |
| DBD, n (%) | 14 (70.0%) | 18 (72.0%) | 15 (88.2%) |  |  |
| UNOS native lung disease category, n (%) |  |  |  |  |  |
| A: obstructive lung disease | 4 (20.0%) | 6 (24.0%) |  |  |  |
| B: pulmonary vascular disease | 0 (0.0%) | 2 (8.0%) |  |  |  |
| C: cystic fibrosis | 12 (60.0%) | 8 (32.0%) |  |  |  |
| D: restrictive lung disease | 4 (20.0%) | 9 (36.0%) |  |  |  |
| CMV serostatus match, n (%) |  |  |  |  |  |
| D-/R- | 5 (25.0%) | 7 (28.0%) |  |  |  |
| R+ | 7 (35.0%) | 13 (52.0%) |  |  |  |
| D+/R- | 7 (35.0%) | 5 (20.0%) |  |  |  |
| Unknown | 1 (5.0%) | 0 (0.0%) |  |  |  |
| Infection**†, n (%) | 12 (60.0%) | 7 (28.0%) | 2 (11.8%) | ND | 2 (12.5%) |
| Bacterial/fungal colonization**, n (%) | 8 (40.0%) | 7 (28.0%) |  |  |  |
| Calcineurin inhibitor**, n (%) |  |  |  |  |  |
| Tacrolimus | 17 (85.0%) | 18 (72.0%) |  |  |  |
| Cyclosporine | 3 (15.0%) | 6 (24.0%) |  |  |  |
| Unknown | 0 (0.0%) | 1 (4.0%) |  |  |  |
| Cell cycle inhibitor**, n (%) |  |  |  |  |  |
| Mycophenolate mofetil | 19 (95.0%) | 18 (72.0%) |  |  |  |
| Azathioprine | 1 (5.0%) | 4 (16.0%) |  |  |  |
| None | 0 (0.0%) | 2 (8.0%) |  |  |  |
| Unknown | 0 (0.0%) | 1 (4.0%) |  |  |  |
| Prednisone dose** |  |  |  |  |  |
| Mean ± SD (mg/day) | 15.4 ± 6.8 | 13.5 ± 5.6 | | | |
| Unknown, n (%) | 0 (0.0%) | 1 (4.0%) |  |  |  |
| Historic DSA, n (%) | 15 (75.0%) | 15 (60.0%) |  |  |  |
| Historic DQ DSA, n (%) | 12 (60.0%) | 12 (48.0%) |  |  |  |
| Current DSA**, n (%) | 7 (35.0%) | 8 (32.0%) |  |  |  |

RLO, restrictive allograft syndrome-like opacities; CLAD, chronic lung allograft dysfunction; IPF, idiopathic pulmonary fibrosis; IQR, interquartile range; SD, standard deviation; R, recipient; D, donor; M, male; F, female; EVLP, ex vivo lung perfusion; DBD, donation after brain death; UNOS, United Network for Organ Sharing; CMV, cytomegalovirus; DSA, donor-specific antibody; ND, no data
\*Donor smoking history for CLAD cases
\*\*At time of explant
†Infection is defined as the presence of a pathogen and the initiation of treatment within 2 weeks prior to explant.

The median age at the time of explant or surgery was 25.5 years (IQR: 19.0–48.0) for the RLO-CLAD group and 33.0 years (IQR: 21.0–46.5) for the non-RLO-CLAD group, whereas patients in the lobectomy lung and IPF groups were older (median 68.0 and 65.5 years, respectively). Female patients accounted for 40.0% of the RLO-CLAD group and 64.0% of the non-RLO-CLAD group. Regarding the primary lung disease, cystic fibrosis was the most common indication in the RLO-CLAD group (60.0%), while the non-RLO-CLAD group showed a more distributed range, including restrictive lung disease (36.0%) and cystic fibrosis (32.0%). At the time of explant, infection was more frequently observed in the RLO-CLAD group than in the non-RLO-CLAD group (60.0% vs. 28.0%, p=0.0389). DSA were detected, at any time post-transplant, in 75.0% of RLO-CLAD and 60.0% of non-RLO-CLAD patients (p=1.000).

### Global gene expression profiling

There were no significant differences in gene expression between central and peripheral CLAD samples in a paired analysis using a false discovery rate threshold of adjusted p-value <0.05 (see Supplemental Figure 1). Based on this information, for subsequent analyses that incorporated only one sample per CLAD patient, we used the peripheral sample when available and otherwise we used the central sample for any given CLAD patient.

The PCA analysis revealed a distinct clustering pattern primarily driven by clinical diagnosis. PC1, which accounted for 19% of the total variance, showed a clear separation of the IPF samples from the other phenotypes. PC2 with 13% variance further delineated the CLAD subtypes; specifically, samples identified as RLO-CLAD tended to cluster toward the positive axis of PC2, whereas non-RLO-CLAD and control samples showed closer proximity and partial overlap (Figure 1). These results confirm that major sources of transcriptomic variation in this cohort are associated with specific disease-related remodeling and clinical phenotypes.

**Figure 1.**
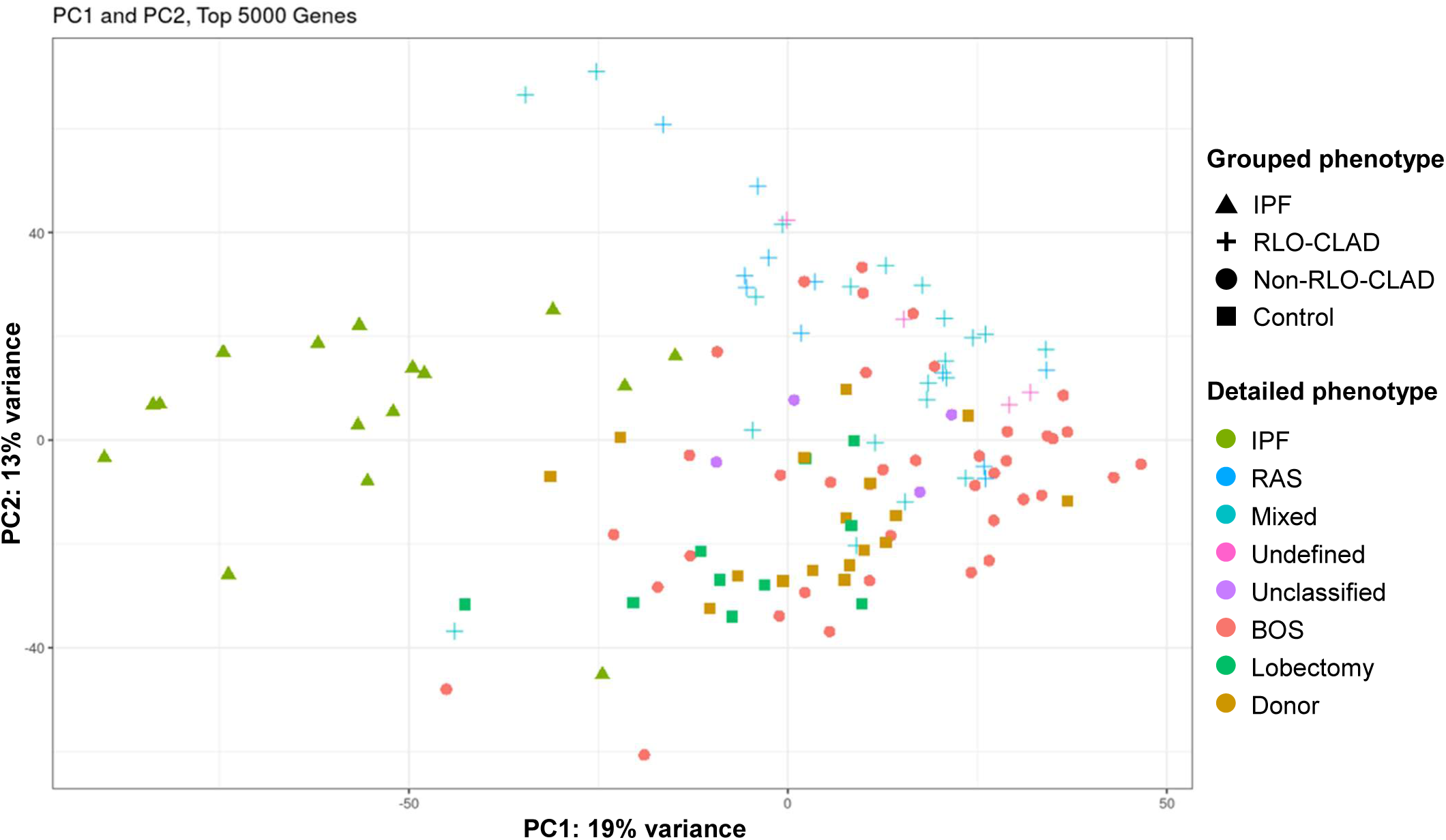
Principal Component Analysis (PCA) of global transcriptomic profiles. PCA based on the top 5000 variable genes demonstrates clustering driven by clinical phenotype. PC1 (19% variance) primarily separates IPF from other groups. PC2 (13% variance) delineates CLAD subtypes, with RLO-CLAD samples clustering toward the positive axis, while non-RLO-CLAD and controls show closer proximity and partial overlap. Symbols represent grouped phenotypes; colors indicate detailed adjudicated phenotypes.

### Identification of Distinct Transcriptomic Signatures via NMF

NMF identified seven distinct gene expression signatures, with the heatmap of the top 50 highest weighted genes for each signature (Figure 2, Supplemental Table 1) revealing clear clustering patterns that align with specific clinical phenotypes. This alignment highlights the capacity of the NMF approach to effectively capture diverse transcriptomic profiles. For select NMF signatures, we compared the overall signature scores between clinical phenotypes (Figure 3). To define the biological functions of these signatures, we conducted pathway enrichment analysis and examined the top 10 ranked genes for each signature (Figure 3). While signature 6 showed no specific pathway enrichment, the remaining six signatures exhibited distinct biological themes. Based on enriched GO terms and key genes, we assigned the following functional labels and list representative top-ranked genes: Signature 0 (epithelial homeostasis; *SCGB3A2, WIF1, SFTA3, FOS, PLA2G4F*), Signature 1 (extracellular matrix remodeling; *COL3A1, COL1A1, COL1A2, POSTN*), Signature 2 (injury response; *SFTPC, SFTPA1, SFTPA2, NAPSA*), Signature 3 (B-cell response; *IGHG2, IGKC, IGLC* family members), Signature 4 (innate immunity; *FCN3, S100A8/A9/A12, NKG7, DEFA1B*), and Signature 5 (cilium/microtubule; *C20orf85 [CIMIP1], C9orf24 [SPMIP9], C1orf194 [CFAP276], FAM183A [CFAP144]*) (Figure 3).

**Figure 2.**
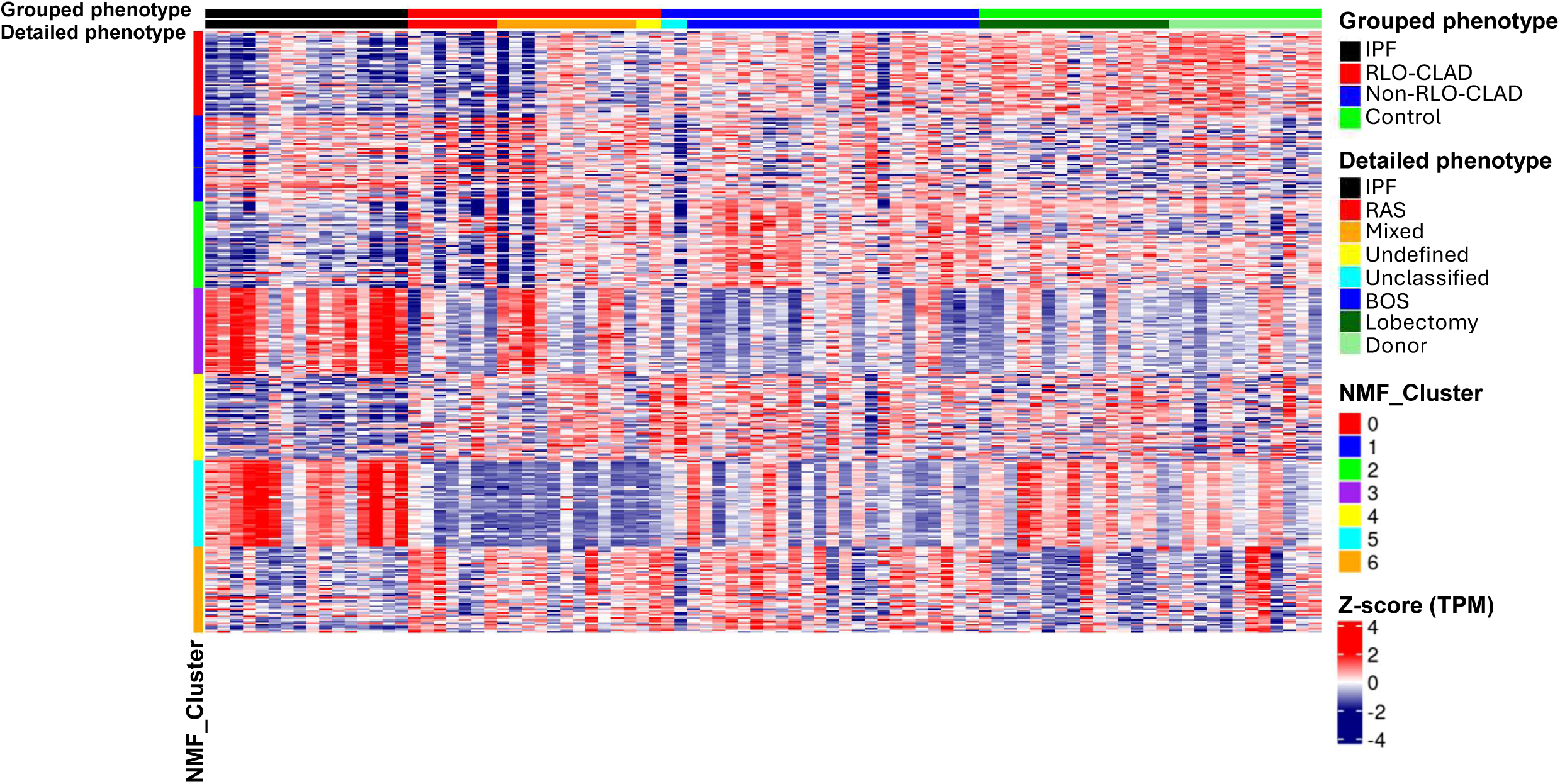
Heatmap of NMF-derived gene expression signatures. Heatmap is displaying the top 50 highly weighted genes for each of the seven signatures identified by non-negative matrix factorization (NMF). Rows represent genes grouped by NMF cluster (Signatures 0–6), and columns represent individual samples. Top annotations indicate the grouped phenotypes and detailed phenotypes. The clustering patterns demonstrate alignment between transcriptomic profiles and clinical phenotypes.

**Figure 3.**
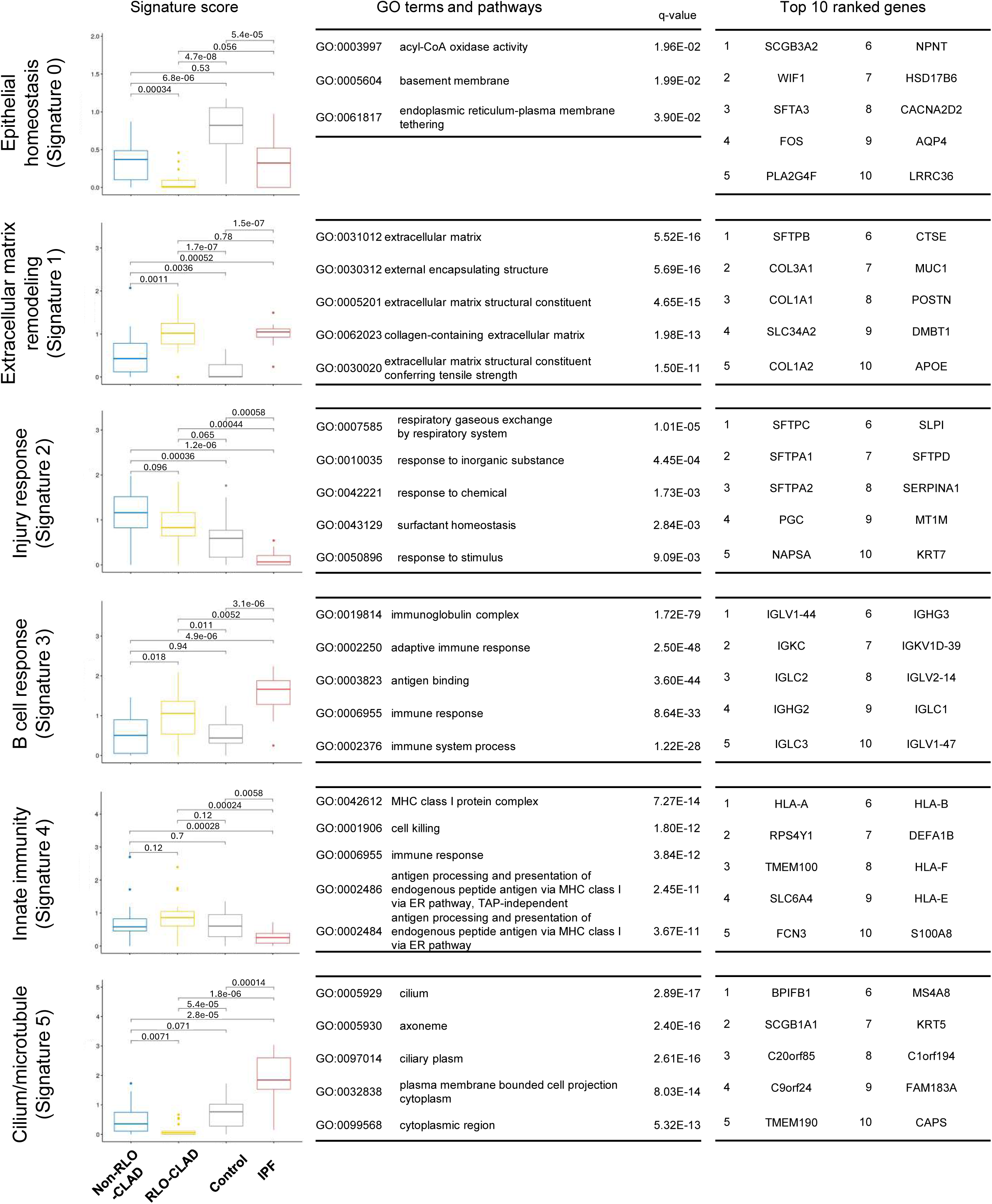
Functional characterization and phenotype association of NMF-derived signatures. This panel illustrates the biological themes and clinical associations of the key transcriptomic signatures identified via non-negative matrix factorization (NMF), displaying signature score distributions across grouped phenotypes, enriched Gene Ontology (GO) terms and pathways with corresponding q-values (representing p-values adjusted for the false discovery rate to account for multiple hypothesis testing), and the top 10 ranked genes contributing to each profile. RLO-CLAD and IPF samples demonstrate significant enrichment of the extracellular matrix remodeling signature (Signature 1), while RLO-CLAD is also uniquely characterized by a B cell response transcriptional footprint (Signature 3). Conversely, signatures associated with epithelial homeostasis (Signature 0) and cilium/microtubule function (Signature 5) show a progressive decline from control lungs to non-RLO-CLAD, with the most profound suppression observed in the RLO-CLAD group. The injury response signature (Signature 2), which includes alveolar type II markers and surfactant-related genes, is relatively enriched in the non-RLO-CLAD phenotype compared to other groups.

### Association Between NMF-Derived Signatures and CLAD Phenotypes

Distinct transcriptomic signatures identified by NMF differentiate RLO-CLAD and non-RLO-CLAD phenotypes (Figure 3). Two signatures—extracellular matrix remodeling and B-cell response—were strongly associated with RLO-CLAD, whereas three signatures—epithelial homeostasis, injury response, and cilium/microtubule—were significantly downregulated in RLO-CLAD compared with non-RLO-CLAD. These differences were confirmed by signature score distributions, which demonstrated clear separation between phenotypes (Figure 3, first column). The extracellular matrix remodeling signature was most prominent in RLO-CLAD and IPF samples, consistent with the fibrotic pathology observed in these groups. The strong enrichment of the B-cell response signature in RLO-CLAD suggests a dominant humoral immune component in restrictive phenotypes.^16^ Among the signatures downregulated in RLO-CLAD, epithelial homeostasis signature gene expression declined progressively from control lungs to non-RLO-CLAD and was lowest in RLO-CLAD, indicating a marked loss of epithelial homeostatic genes in RLO-CLAD. The injury response signature was positively correlated with non-RLO-CLAD, showed intermediate levels in RLO-CLAD, and was lowest in IPF. Expression of the cilium/microtubule signature was markedly reduced in RLO-CLAD compared to non-RLO-CLAD and control lungs. A volcano plot comparing gene expression profiles between RLO-CLAD and non-RLO-CLAD is presented in Supplemental Figure 2, with the top 10 genes selected from the top 50 ranked genes for each signature highlighted,

### Clinical Correlates of Transcriptomic Signatures

The clinical associations of the NMF transcriptomic signatures are detailed in Figure 4. Both RLO-CLAD and lungs with pathological evidence of RAS-like features demonstrated strong positive correlations with extracellular remodeling and B-cell response signatures, along with negative correlations with epithelial homeostasis and cilium/microtubule signatures. Similarly, radiological RLO were specifically and strongly associated with the extracellular remodeling signature. In contrast, radiological air trapping or hyperinflation showed a strong positive correlation with the injury response signature but were negatively correlated with extracellular remodeling and innate immunity. Female recipients exhibited weak positive correlations with injury response and cilium/microtubule signatures, and weak negative correlations with B-cell and innate immune responses. CMV serologic mismatch between donor and recipient was associated with B-cell and innate immunity, whereas it correlated negatively with the injury response signature. Additionally, the presence of infection at the time of explant was associated with increased extracellular remodeling and decreased epithelial homeostasis. Finally, a history of DSA, any time prior to explant, was strongly associated with the injury response signature, while showing negative correlations with epithelial homeostasis and cilium/microtubule signatures. Additional associations of all NMF signatures and clinical parameters are provided in Supplemental Figure 3.

**Figure 4.**
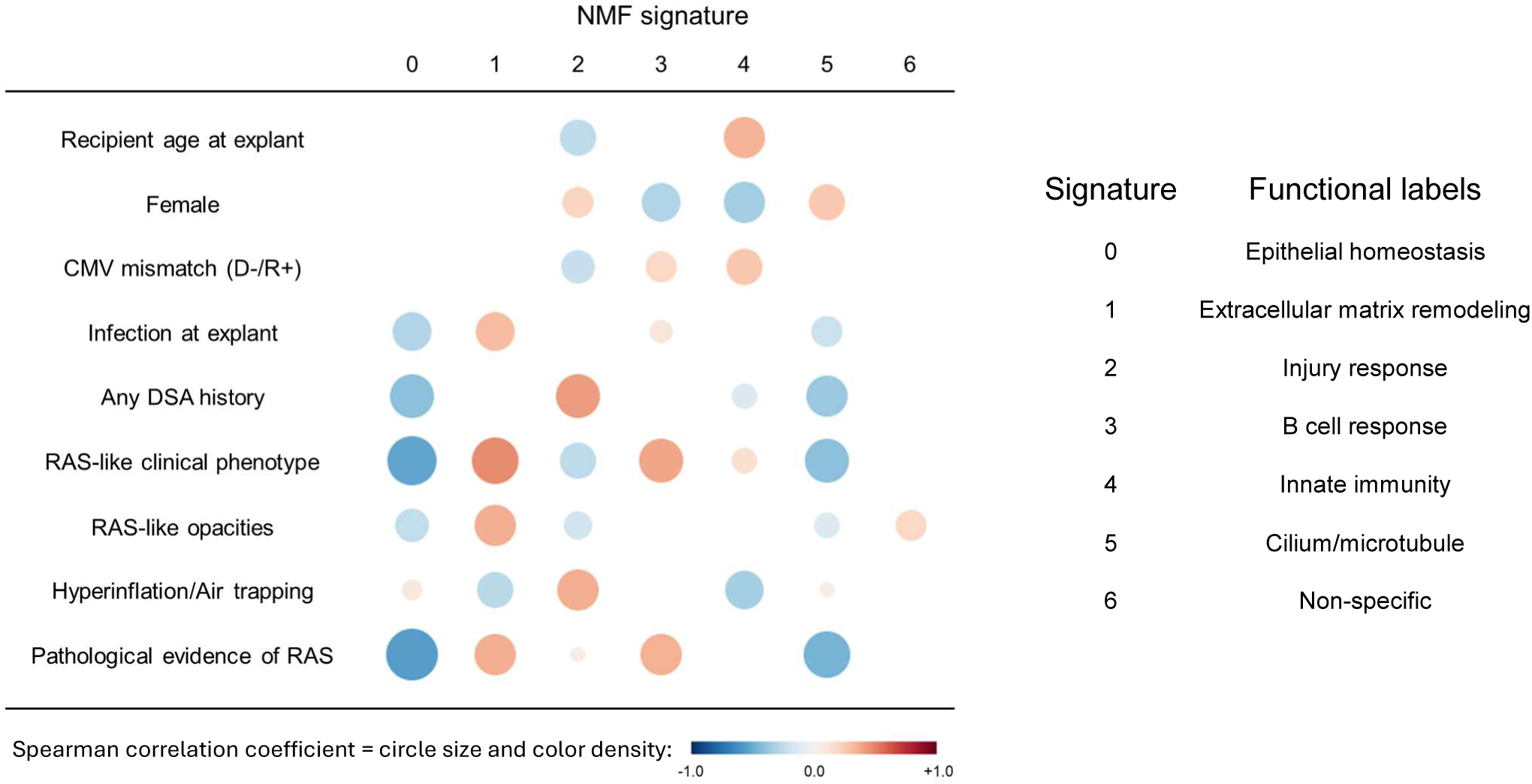
Correlation of NMF signatures with clinical and pathological variables. Dot plot showing Spearman correlations between the seven NMF-derived signatures and clinical parameters. Circle sizes and color densities represent the strength of the clinical associations, scaled according to the Spearman correlation coefficient (red: positive; blue: negative). RAS-like clinical and pathological phenotypes correlate positively with extracellular matrix remodeling (Signature 1) and B cell response (Signature 3), while correlating negatively with epithelial homeostasis (Signature 0) and ciliary (Signature 5) signatures. Hyperinflation/air trapping is positively associated with the injury response (Signature 2). DSA history and infection also exhibit distinct signature-specific correlation patterns.

### SCGB3A2 protein levels in bronchoalveolar lavage

SCGB3A2 was among the most highly ranked genes within the epithelial homeostasis signature, which correlated negatively with RLO-CLAD. BAL analysis demonstrated significantly lower SCGB3A2 concentrations in CLAD compared to non-CLAD controls (p<0.001). Although there was a trend towards lower SCGB3A2 in RLO-CLAD compared to non-RLO-CLAD, this difference was not statistically significant (p=0.080) (Figure 5).

**Figure 5.**
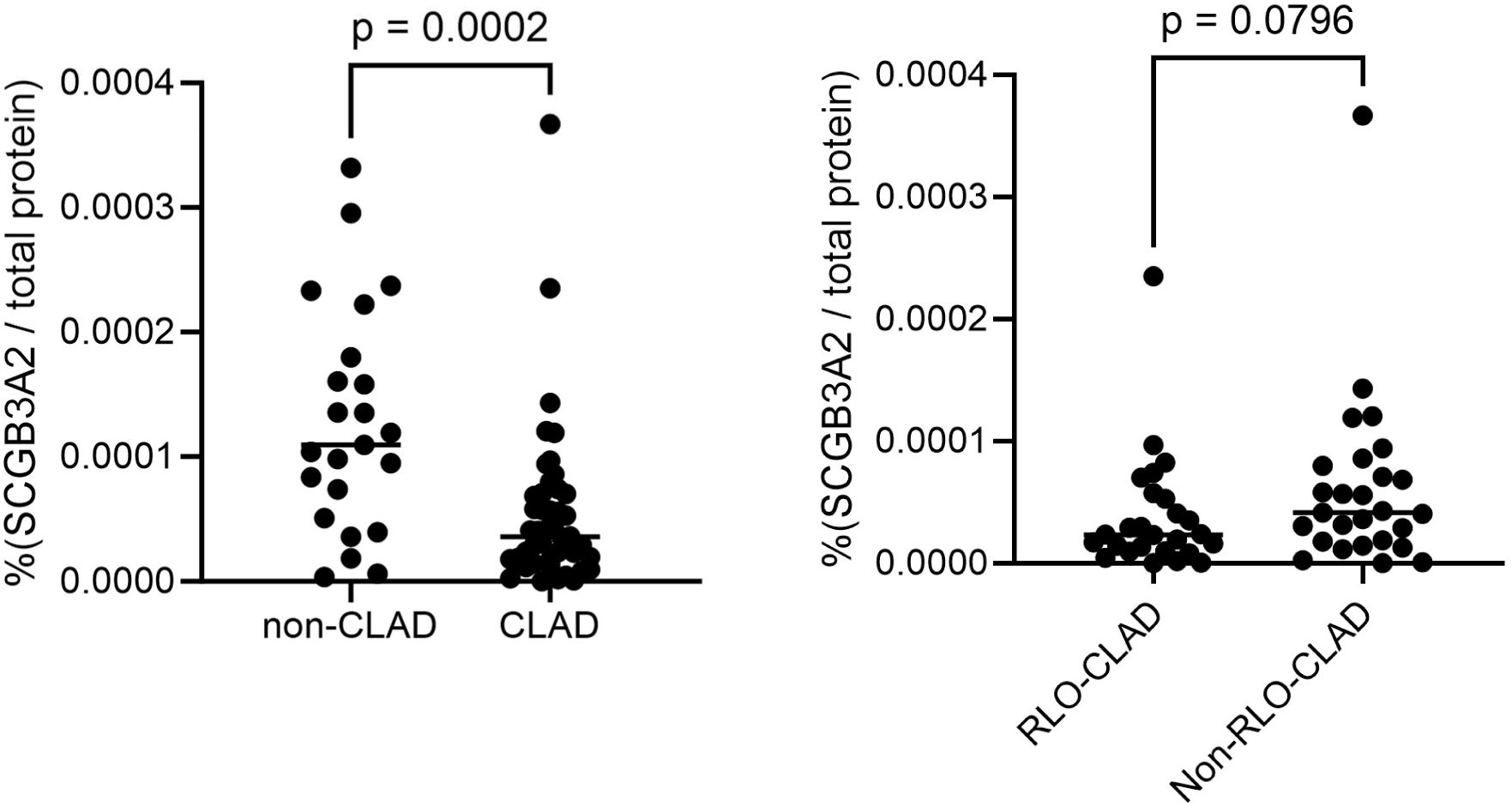
SCGB3A2 protein levels in bronchoalveolar lavage (BAL) fluid. SCGB3A2 concentrations normalized to total protein were measured in an independent cohort of lung transplant recipients. BAL SCGB3A2 levels are significantly reduced in the CLAD group compared to non-CLAD controls (left). While there is a downward trend in RLO-CLAD compared to non-RLO-CLAD, the difference did not reach statistical significance (right). Comparisons between groups were performed using the Mann-Whitney test. p-values are indicated for each comparison.

### Ciliated cell counts by immunofluorescence staining

Since the cilium/microtubule signature was reduced in RLO-CLAD compared to non-RLO-CLAD, we assessed ciliated cells by FOXJ1⁺ immunofluorescence staining. Although quantitative analysis showed trends toward lower proportions of FOXJ1⁺ ciliated epithelial cells in CLAD, particularly in RLO-CLAD, these differences were not statistically significant (Figure 6). Together, these findings suggest that suppression of ciliary transcriptional programs in RLO-CLAD may reflect functional epithelial dysregulation rather than overt depletion of ciliated cells.

**Figure 6.**
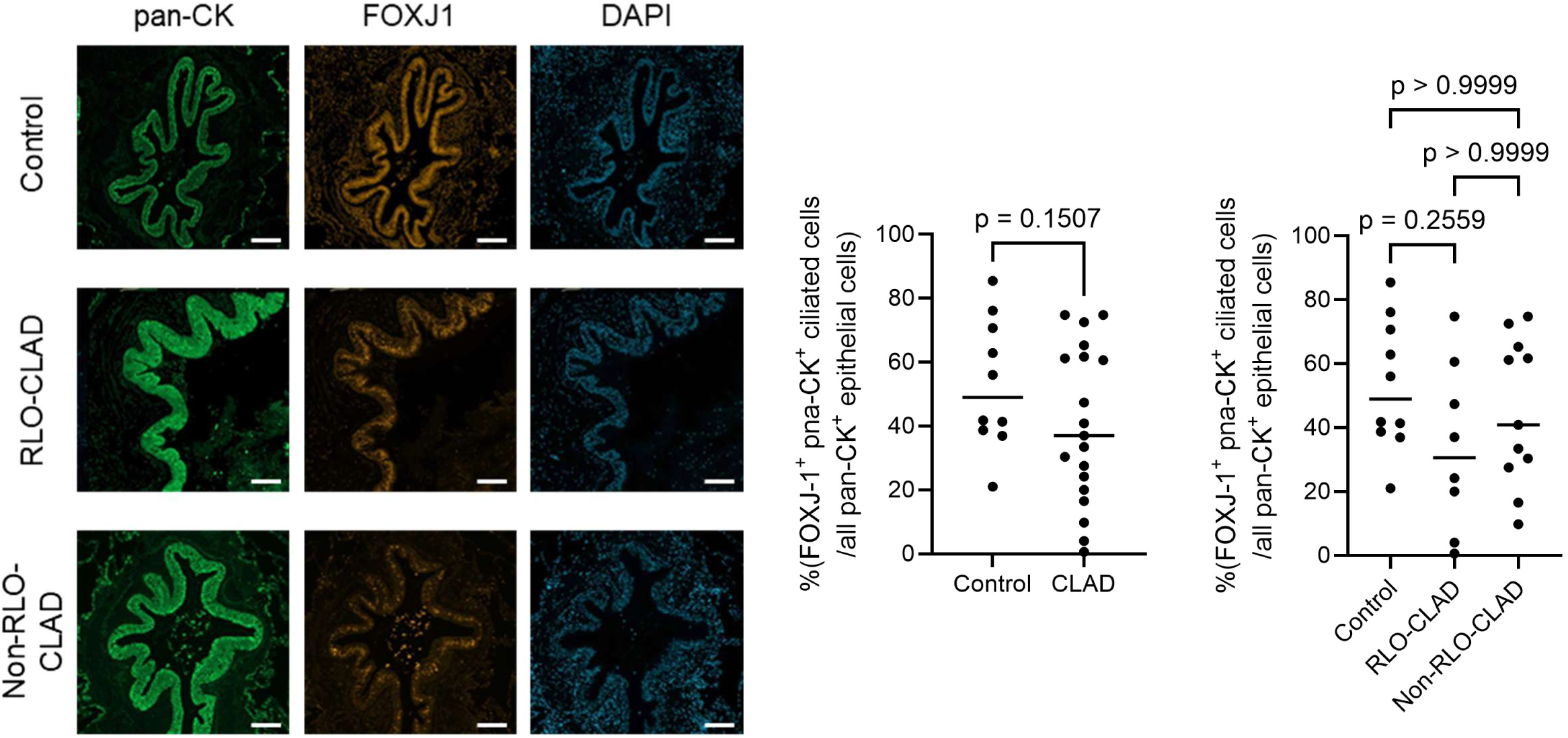
Assessment of airway ciliated cell composition. Representative immunofluorescence images (left) show staining for pan-cytokeratin (pan-CK; green), the ciliated cell marker FOXJ1 (orange), and DAPI (blue) in lung tissue from control, RLO-CLAD, and non-RLO-CLAD groups. Scale bar = 100 µm. Quantitative analysis (right) of the percentage of FOXJ1^+^, pan-CK^+^ ciliated cells among all pan-CK^+^ epithelial cells revealed no statistically-significant differences between phenotypes. While a trend towards lower ciliated cell proportions was observed in RLO-CLAD, these findings may suggest that the marked transcriptional repression of ciliary genes reflects functional dysregulation rather than an overt loss of cells.

## Discussion

This study provides a comprehensive transcriptomic stratification of RLO-CLAD and non-RLO-CLAD using explanted CLAD lungs, revealing distinct gene signatures that distinguish their divergent clinical phenotypes. By leveraging NMF, an unsupervised dimensionality-reduction and feature-extraction method, we identified principal themes that emerge from the data: (1) RLO-CLAD is characterised by a prominent profibrotic signature that parallels IPF, (2) RLO-CLAD shows a robust humoral-immune (B cell/plasma cell) transcriptional footprint, (3) loss of epithelial homeostatic signature is a shared feature across CLAD but is most prominent in RLO-CLAD, and (4) a downregulation of ciliary and microtubule-associated genes is demonstrated, suggesting profound epithelial dysfunction in RLO-CLAD.

The enrichment of extracellular matrix remodeling genes in RLO-CLAD underscores its well-established clinicopathologic identity as a parenchymal fibrotic process, typically characterized by pleuroparenchymal fibroelastosis and diffuse alveolar fibrosis on histology and imaging. Our findings that IPF explants share aspects of the extracellular matrix remodeling signature with RAS is concordant with prior reports suggesting molecular overlap between fibrotic forms of CLAD and primary fibrotic lung disease.^17^ The leading genes in this pro-fibrotic signature in CLAD in our dataset (e.g., *COL3A1, COL1A1, POSTN, MMP7*) mirror processes in lung fibrosis, indicating that RLO-CLAD exhibits a transcriptional signature comparable to advanced interstitial lung disease. *THY1*, which showed a high signature score, was expressed at approximately four-fold higher levels in RLO-CLAD than in control lungs. Previous studies have also shown that myofibroblasts/fibroblasts express high levels of the THY1 cell-surface antigen in the fibrotic pleura of both the pleuroparenchymal fibroelastosis model in mice and in patients with pleuroparenchymal fibroelastosis.^18^ This signature is also characterized by genes that are associated with epithelial dysfunction and remodeling (e.g., *SFTPB, MUC1*) and their alteration has been previously described in fibrotic lung diseases,^19^ and CLAD.^20^ Collectively, these findings suggest that therapies targeting fibrogenesis, particularly within shared downstream fibrotic effector pathways, could be conceptually relevant to RLO-CLAD, including RAS and mixed phenotypes. Nevertheless, since the present data are cross-sectional and derived from end-stage explants, they cannot determine whether fibrotic programming represents an initiating event in RLO-CLAD or a late, self-sustaining state following diverse injuries. Additionally, it is important to note that IPF and RAS-like pathology are usually quite different, with distinct distribution of fibroblasts and extracellular matrix: these differences should be a major focus of future studies that should incorporate a spatial component to these analyses. Such studies may help glean which therapies may be extrapolatable from IPF to CLAD in the future.

The prominent upregulation of B cell response-associated transcripts in RLO-CLAD, captured in Signature 3 (e.g., *IGKC, IGLC2, IGHG3, IGLV1-47, MZB1*), is consistent with the recognized role of humoral immunity in restrictive graft failure.^5^ Although a causative role of B cells in RLO-CLAD is not entirely confirmed, immunohistochemical analyses of RLO-CLAD explants have identified B-cell lymphoid structures or follicles predominantly in RLO-CLAD, with lesser involvement in non-RLO-CLAD or control lungs.^16^ A mouse model of RAS-like pathology similarly demonstrated increased plasma cell infiltration, and plasmacytic vasculitis in lungs.^21,22^ Notably, a recent study using two different RAS mouse models reported that bortezomib and rituximab depleted antibody-secreting cells and attenuated fibrosis,^23^ and B cell depletion decreased RAS-like fibrosis.^21^ Our findings reinforce these observations and suggest that B cell–targeted therapies may warrant phenotype- or endotype-specific trials. Taken together, these data support the biological plausibility of B cell-mediated pathways as a major driver of RLO-CLAD / RAS. Human case series have shown associations between circulating DSA, antibody-mediated rejection, and development of RAS pathology.^24–26^ Multiple clinical studies link DSA to CLAD and adverse outcomes, but their distribution between RAS and BOS varies across cohorts: some studies identify DSA in both RAS and BOS,^27,28^ while others implicate HLA class II antibodies in RAS specifically,^29^ and still others emphasize *de novo* DSA as a risk factor for BOS progression.^30–32^ While our data underscore a notable humoral signature in RLO-CLAD, our NMF analysis showed that a history of serum DSA was not more strongly associated with the B cell response signature than with epithelial injury response signature. As previously reported, DSA in lung tissue correlates with RAS compared to BOS, whereas serum DSA does not strongly discriminate between phenotypes.^26^ These variabilities may reflect differences in timing (early vs late DSA), specificity (class I vs class II), sample type (serum vs tissue), assay sensitivity, clinical management practices (surveillance intensity, treatment of DSA), and the presence of confounding variables such as infections.

In contrast, the epithelial homeostasis gene signature (Signature 0) was diminished across CLAD, particularly in RLO-CLAD (control > non-RLO-CLAD > RLO-CLAD). SCGB3A2, a secretoglobin with epithelial protective functions and predominant expression in airway epithelial club cells,^33^ ranked most prominently within this signature and showed marked downregulation in RLO-CLAD. SCGB3A2 has been reported to exert anti-inflammatory and anti-fibrotic effects. In an ovalbumin-induced airway inflammation mouse model, reduced pulmonary *SCGB3A2* transcript levels were inversely correlated with higher concentrations of proinflammatory cytokines in BAL fluid.^34–36^ In a bleomycin-induced pulmonary fibrosis mouse model, SCGB3A2 also attenuated the expression of fibrosis-related genes by inhibiting TGF-β signaling, thereby limiting myofibroblast differentiation.^37^ Several studies, including a recent multicenter study, have demonstrated that concentrations of another secretoglobin, club cell secretory protein (CCSP; encoded by *SCGB1A1*), in BAL fluid were associated with concurrent CLAD and future CLAD risk, suggesting its utility for early post-transplant risk stratification.^38,39^ To the best of our knowledge, the relationship between SCGB3A2 and CLAD risk has not been investigated to date. In our study, BAL protein validation demonstrated lower SCGB3A2 protein levels at CLAD onset compared with non-CLAD samples, supporting its potential utility as an early detection biomarker. Functionally, club cells contribute to airway epithelial repair, immune modulation, and secretion of protective factors.^40,41^ Their depletion plausibly compromises mucosal defense and reparative capacity, facilitating both airway obliteration in BOS and aberrant fibroproliferation in RAS. Notably, the transcriptomic decline of *SCGB3A2* gene in RLO-CLAD was significantly more pronounced than in non-RLO-CLAD in our explant tissues, whereas BAL SCGB3A2 levels were reduced in CLAD overall with only a non-significant trend towards reduction in RLO-CLAD compared to non-RLO-CLAD. Several reasons may explain this discrepancy. First, spatial heterogeneity of club cell loss (e.g., more severe peripheral airway involvement in regions contributing to RAS pathology) may lead to a more pronounced apparent reduction in tissue transcripts without a proportional change in more centrally sampled BAL. Second, post-transcriptional regulation, altered secretion, or protein degradation may affect BAL protein levels differently from tissue transcripts. Third, differences in sampling timing may affect the concordance between explant transcriptomics and BAL protein levels at CLAD onset. These possibilities underscore the need for spatially resolved single-cell and proteomic studies to map club cell loss and dysfunction across airway compartments and disease stages. Whether restoration of secretoglobin function can mitigate chronic allograft injury remains an important question for future investigation.

Surfactant/injury response gene signatures (Signature 2) were upregulated in CLAD, and highest in non-RLO-CLAD. This signature contained alveolar type II markers (*SFTPC, SFTPA, NAPSA*). Surfactant gene upregulation is consistent with an ongoing epithelial stress response and suggests that non-RLO-CLAD may involve more active alveolar epithelial remodeling than RLO-CLAD. The association between Signature 2 and DSA raises the possibility that humoral injury contributes to epithelial stress in non-RLO-CLAD, although causality cannot be inferred. This pattern (non-RLO-CLAD > RLO-CLAD > IPF) suggests differing dynamics of epithelial injury and repair in each phenotype.

The downregulation of ciliary and microtubule-associated genes (Signature 5) in RLO-CLAD indicates substantial airway epithelial dysfunction in addition to its characteristic fibrotic pathology. Key components of mucociliary clearance, indicating *FOXJ1*, a transcription factor essential for ciliogenesis,^42,43^ and multiple genes in its family, were among the most markedly suppressed in RLO-CLAD. Immunohistochemistry showed no significant differences in FOXJ1⁺ ciliated cell counts among groups, implying transcriptional dysregulation or altered differentiation rather than overt loss of ciliated cells. There was a mild trend towards reduced FOXJ1^+^ cells in CLAD compared to control, suggesting that a subset of patients may have particularly reduced ciliated cells: further exploring this endotype may identify patients with greater epithelial injury and altered physiology. Prior studies in chronic airway diseases show that ciliary dysfunction, manifesting as reduced beat frequency or dyskinetic motion, can occur despite preserved ciliated cell numbers.^44,45^ Therefore, the transcriptional reductions we observed could reflect functional impairment not captured by cell quantitation. Functional assays of mucociliary clearance and high-resolution imaging of ciliary ultrastructure would clarify whether ciliary gene repression contributes to disease mechanisms and phenotype differences. Whether these transcriptional changes represent primary pathogenic drivers or secondary remodeling responses remains unclear, but they highlight epithelial integrity as a potential therapeutic target.

It is worth noting that while our NMF analysis clustered genes into signatures that had clear correlations with RLO-CLAD and non-RLO-CLAD phenotypes, the distribution observed in our PCA suggests a continuous molecular spectrum rather than completely segregated, binary endotypes. This aligns with a debate over whether clinical CLAD phenotypes represent distinct biologic entities or a continuum of molecular changes driven by advancing stages of tissue fibrosis and fibroproliferative remodeling, with disparate spatial involvement. Future longitudinal studies tracking molecular kinetics from CLAD onset to explant are needed to clarify this distinction.

There are several limitations to this study. First, bulk RNA sequencing captures averaged gene expression across heterogeneous cell populations, limiting the ability to resolve specific cellular contributors. Although NMF allowed the identification of coherent gene signature patterns, the biological interpretation of these signatures relies on gene ranking and pathway enrichment and therefore requires further validation. Moreover, the associations identified between signatures and clinical variables do not establish causation. These findings should be validated in larger, longitudinal studies incorporating single-cell analyses. Second, differences in timing and sampling warrant consideration. This study is based on explanted lungs from patients with end-stage CLAD, and the findings may not fully represent earlier disease stages. The BAL cohort was independent of the explant cohort, and samples were obtained at the time of CLAD onset. In addition, potential differences in sampling conditions may have influenced protein level measurements. Immunofluorescence analysis was limited to selected tissue regions and may not reflect global epithelial distribution. Third, the results should be interpreted with caution given the heterogeneity in immunologic exposures and clinical management across centers. Immunosuppressive regimens, institutional practices regarding DSA surveillance and treatment, and co-morbid infections, particularly CMV, vary across cohorts and can modulate immune activation and repair responses. These factors plausibly shift the relative expression of immune, epithelial, and fibrotic transcripts.

## Conclusions

The present study, using NMF analysis, delineates tissue-level transcriptomic patterns demonstrating that RLO-CLAD and non-RLO-CLAD exhibit differential molecular presentations within CLAD. The RLO-CLAD group is characterized by more prominent profibrotic and B cell response signatures and by downregulated ciliary and microtubule-associated signature; whereas, non-RLO-CLAD shows a relative enrichment of alveolar epithelial injury–response pathways compared with RLO-CLAD. Epithelial homeostasis, including secretoglobin expression, appears to represent a shared vulnerability across CLAD, with particularly pronounced reduction in RLO-CLAD. Rather than representing strict binary endotypes, these signatures likely reflect a continuum of molecular reprogramming driven by the severity and location of fibroproliferative processes. These observations provide novel insights into CLAD heterogeneity that warrant further investigation in longitudinal and mechanistic studies.

## Acknowledgements

The authors would like to thank Dr. Shane Shapera for help with IPF patient identification, Dr. Clara Grizales for help with patient phenotyping, the Toronto Lung Transplant Program (TLTP) database team and the TLTP biobank team. We thank the many laboratory members who participated in data collection and sample processing, particularly those who processed explanted CLAD lung tissue for bulk RNA sequencing: Hisashi Oishi, Jussi Tikkanen, Yohei Taniguchi, Alex Tigert, Liran Levy, Ei Miyamoto, Allen Duong, Matthew White, Kristen Boonstra, Madalina Maxim, Mitsuaki Kawashima, Akihiro Takahashi, Tina Daigneault, Christina Lee, Goodness Madu, Sajad Moshkelgosha, Benjamin Renaud-Picard, Nam Topp-Nguyen, Gafoor Puthiyaveetil, Daniel Vosoughi, Nadia Sachewsky, and Chen Yang (Kevin) Zhang.

---

Supplemental Figure 1 Volcano plot showing differential expression analysis between central and peripheral CLAD samples. The x-axis represents the log2 fold change, and the y-axis represents the –log10 p-value. The vertical red lines indicate the log2 fold change thresholds (±1), and the horizontal red line represents the adjusted p-value threshold for statistical significance (adjusted p=0.05). No genes reached the threshold for statistical significance, confirming the transcriptomic similarity between the two regions.

Supplemental Figure 2 Differential gene expression between RLO-CLAD and non-RLO-CLAD. A volcano plot illustrates the global transcriptional differences between RLO-CLAD and non-RLO-CLAD samples. The x-axis represents the log2 fold change (positive values indicating higher expression in RLO-CLAD), and the y-axis represents the –log10 p-value. Horizontal and vertical dashed lines indicate the thresholds for statistical significance (p<0.05) and fold change (|log2FC|>0.5), respectively. Labeled points highlight the top 10 genes selected from the top 50 highest-weighted genes for each of the transcriptomic signatures identified by NMF, demonstrating their distribution across the differential expression profile.

Supplemental Figure 3 Correlation between NMF signatures and extended clinical and demographic parameters. Dot plot showing Spearman correlations between the seven NMF-derived signatures and an extended subset of clinical variables. Circle sizes and color densities represent the strength of the clinical associations, scaled according to the Spearman correlation coefficient (red: positive; blue: negative).

Supplemental Table 1. List of the top 50 highest weighted genes for each of the seven transcriptomic signatures identified by NMF

