## Supplemental figure 1,2,3 for "Distinct fibrotic, epithelial and immune transcriptomic programs in phenotypes of chronic lung allograft dysfunction"

Supplemental Fig. 1

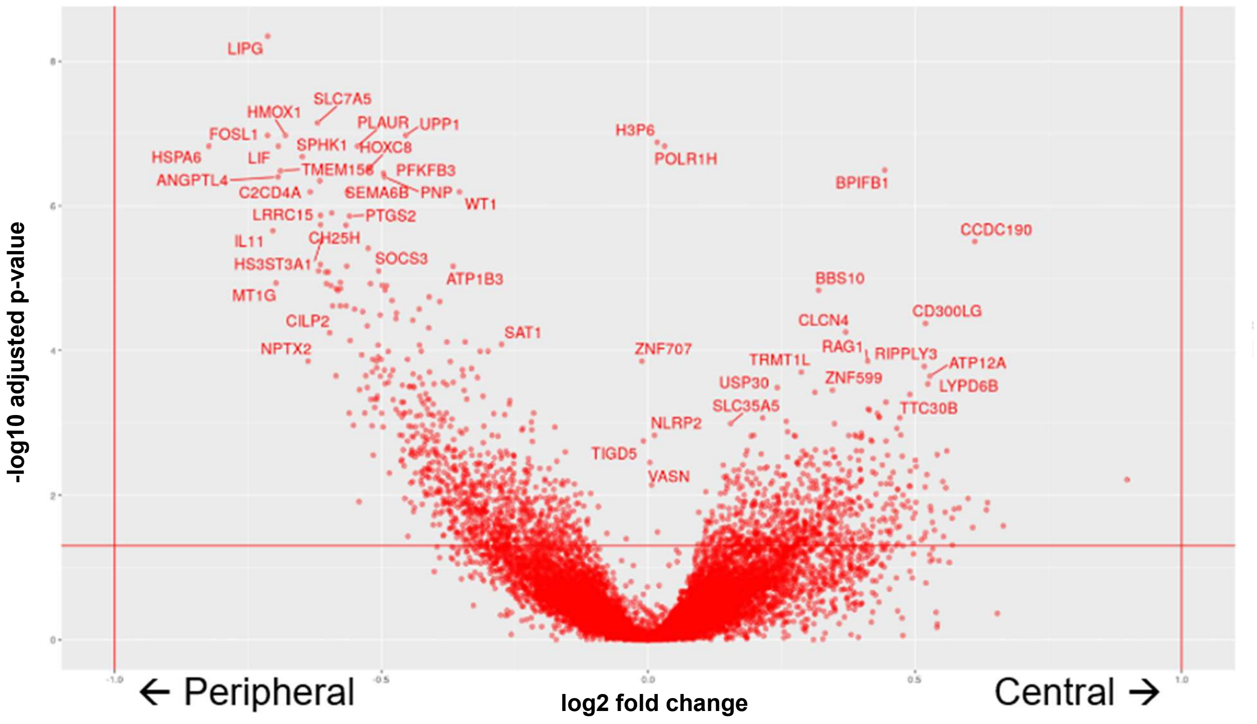


Supplemental Fig. 2

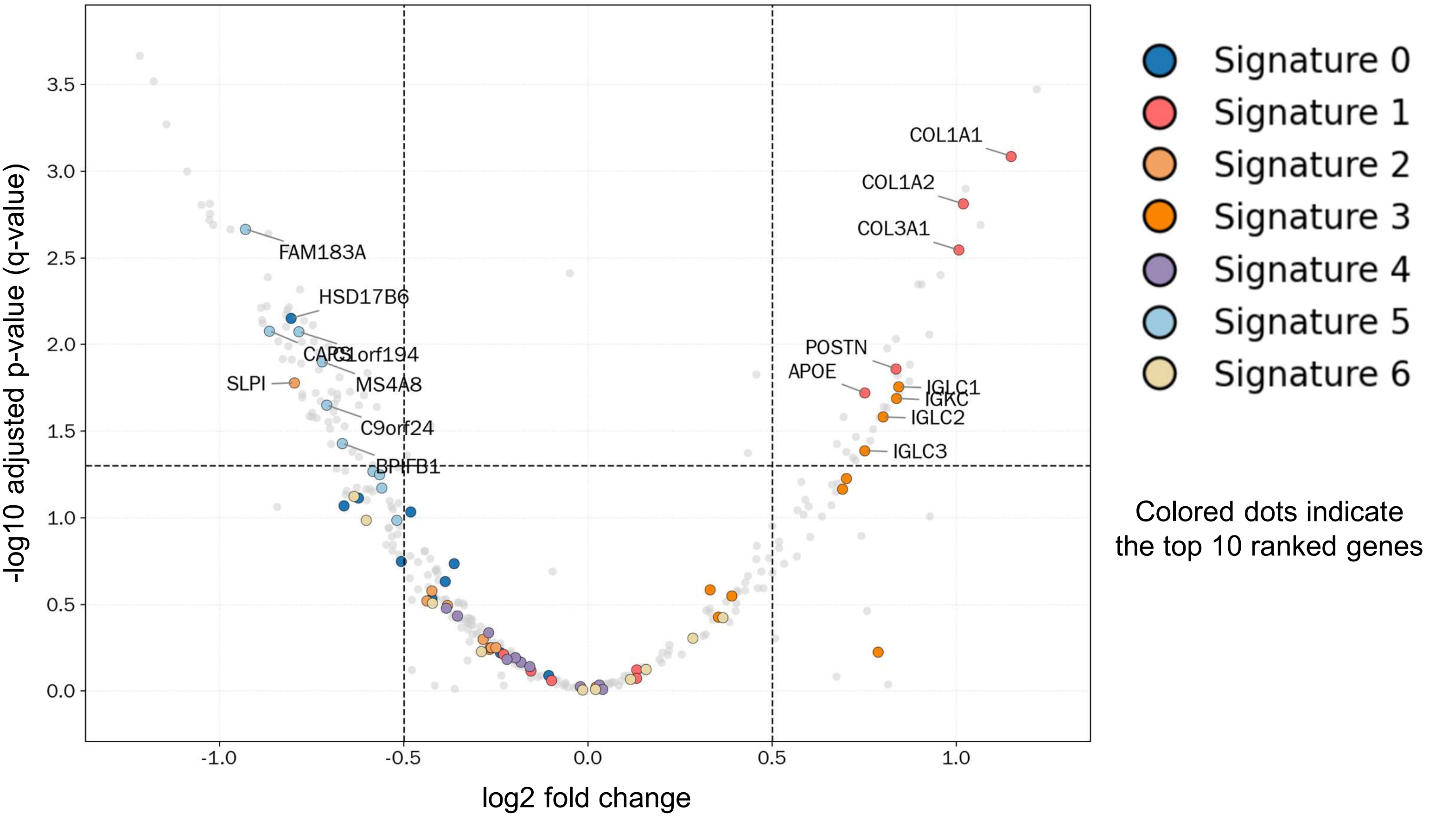


RLO-CLAD vs. non-RLO-CLAD

### Supplemental Fig. 3

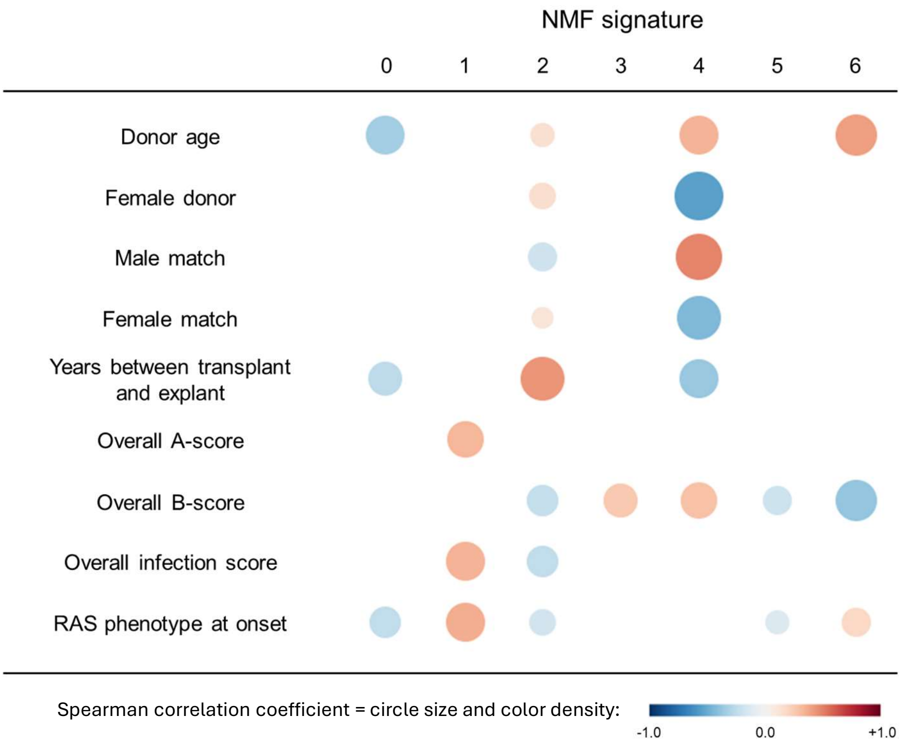


| Signature | Functional labels |
|---|---|
| 0 | Epithelial homeostasis |
| 1 | Extracellular matrix remodeling |
| 2 | Injury response |
| 3 | B cell response |
| 4 | Innate immunity |
| 5 | Cilium/microtubule |
| 6 | Non-specific |
