## Supplemental Table 1 for "Distinct fibrotic, epithelial and immune transcriptomic programs in phenotypes of chronic lung allograft dysfunction"

List of the top 50 highest weighted genes for each of the seven transcriptomic signatures identified by NMF

| Signature | Rank | Gene ID | Gene Name | Signature Score |
|---|---|---|---|---|
| 0 | 1 | ENSG00000164265 | SCGB3A2 | 2.73 |
| 0 | 2 | ENSG00000156076 | WIF1 | 2.29 |
| 0 | 3 | ENSG00000257520 | SFTA3 | 2.24 |
| 0 | 4 | ENSG00000170345 | FOS | 2.2 |
| 0 | 5 | ENSG00000168907 | PLA2G4F | 2.04 |
| 0 | 6 | ENSG00000168743 | NPNT | 2.03 |
| 0 | 7 | ENSG00000025423 | HSD17B6 | 2.03 |
| 0 | 8 | ENSG00000007402 | CACNA2D2 | 1.99 |
| 0 | 9 | ENSG00000171885 | AQP4 | 1.93 |
| 0 | 10 | ENSG00000159708 | LRRC36 | 1.82 |
| 0 | 11 | ENSG00000115361 | ACADL | 1.79 |
| 0 | 12 | ENSG00000104267 | CA2 | 1.78 |
| 0 | 13 | ENSG00000166348 | USP54 | 1.78 |
| 0 | 14 | ENSG00000144331 | ZNF385B | 1.76 |
| 0 | 15 | ENSG00000154065 | ANKRD29 | 1.73 |
| 0 | 16 | ENSG00000169031 | COL4A3 | 1.71 |
| 0 | 17 | ENSG00000197415 | VEPH1 | 1.7 |
| 0 | 18 | ENSG00000112972 | HMGCS1 | 1.69 |
| 0 | 19 | ENSG00000152527 | PLEKHH2 | 1.65 |
| 0 | 20 | ENSG00000160867 | FGFR4 | 1.65 |
| 0 | 21 | ENSG00000053747 | LAMA3 | 1.63 |
| 0 | 22 | ENSG00000175318 | GRAMD2A | 1.5 |
| 0 | 23 | ENSG00000145147 | SLIT2 | 1.5 |
| 0 | 24 | ENSG00000150625 | GPM6A | 1.48 |
| 0 | 25 | ENSG00000158220 | ESYT3 | 1.48 |
| 0 | 26 | ENSG00000165195 | PIGA | 1.46 |
| 0 | 27 | ENSG00000196549 | MME | 1.46 |
| 0 | 28 | ENSG00000135917 | SLC19A3 | 1.41 |
| 0 | 29 | ENSG00000153093 | ACOXL | 1.41 |
| 0 | 30 | ENSG00000189367 | KIAA0408 | 1.38 |
| 0 | 31 | ENSG00000152580 | IGSF10 | 1.36 |
| 0 | 32 | ENSG00000144218 | AFF3 | 1.32 |
| 0 | 33 | ENSG00000206077 | ZDHHC11B | 1.31 |
| 0 | 34 | ENSG00000266524 | GDF10 | 1.29 |
| 0 | 35 | ENSG00000154319 | FAM167A | 1.26 |
| 0 | 36 | ENSG00000167074 | TEF | 1.25 |
| 0 | 37 | ENSG00000173930 | SLCO4C1 | 1.25 |
| 0 | 38 | ENSG00000134216 | CHIA | 1.25 |
| 0 | 39 | ENSG00000176771 | NCKAP5 | 1.22 |
| 0 | 40 | ENSG00000142178 | SIK1 | 1.22 |
| 0 | 41 | ENSG00000103044 | HAS3 | 1.22 |
| 0 | 42 | ENSG00000137460 | FHDC1 | 1.22 |
| 0 | 43 | ENSG00000130518 | IQCN | 1.2 |
| 0 | 44 | ENSG00000088926 | F11 | 1.17 |

| | | | | |
|---|---|---|---|---|
| 0 | 45 | ENSG00000143365 | RORC | 1.15 |
| 0 | 46 | ENSG00000177494 | ZBED2 | 1.14 |
| 0 | 47 | ENSG00000155511 | GRIA1 | 1.12 |
| 0 | 48 | ENSG00000143624 | INTS3 | 1.11 |
| 0 | 49 | ENSG00000255330 | AL096711.2 | 1.1 |
| 0 | 50 | ENSG00000123119 | NECAB1 | 1.09 |
| 1 | 1 | ENSG00000168878 | SFTPB | 3.69 |
| 1 | 2 | ENSG00000168542 | COL3A1 | 3.44 |
| 1 | 3 | ENSG00000108821 | COL1A1 | 3.2 |
| 1 | 4 | ENSG00000157765 | SLC34A2 | 2.93 |
| 1 | 5 | ENSG00000164692 | COL1A2 | 2.9 |
| 1 | 6 | ENSG00000196188 | CTSE | 2.88 |
| 1 | 7 | ENSG00000185499 | MUC1 | 2.76 |
| 1 | 8 | ENSG00000133110 | POSTN | 2.69 |
| 1 | 9 | ENSG00000187908 | DMBT1 | 2.68 |
| 1 | 10 | ENSG00000130203 | APOE | 2.67 |
| 1 | 11 | ENSG00000163359 | COL6A3 | 2.64 |
| 1 | 12 | ENSG00000259803 | SLC22A31 | 2.54 |
| 1 | 13 | ENSG00000154096 | THY1 | 2.52 |
| 1 | 14 | ENSG00000137857 | DUOX1 | 2.52 |
| 1 | 15 | ENSG00000177575 | CD163 | 2.51 |
| 1 | 16 | ENSG00000243716 | NPIPB5 | 2.47 |
| 1 | 17 | ENSG00000130513 | GDF15 | 2.42 |
| 1 | 18 | ENSG00000203697 | CAPN8 | 2.42 |
| 1 | 19 | ENSG00000133048 | CHI3L1 | 2.39 |
| 1 | 20 | ENSG00000137673 | MMP7 | 2.39 |
| 1 | 21 | ENSG00000153395 | LPCAT1 | 2.38 |
| 1 | 22 | ENSG00000101443 | WFDC2 | 2.35 |
| 1 | 23 | ENSG00000164932 | CTHRC1 | 2.34 |
| 1 | 24 | ENSG00000123838 | C4BPA | 2.31 |
| 1 | 25 | ENSG00000188906 | LRRK2 | 2.29 |
| 1 | 26 | ENSG00000185864 | NPIPB4 | 2.28 |
| 1 | 27 | ENSG00000186340 | THBS2 | 2.25 |
| 1 | 28 | ENSG00000169710 | FASN | 2.23 |
| 1 | 29 | ENSG00000047936 | ROS1 | 2.23 |
| 1 | 30 | ENSG00000107796 | ACTA2 | 2.23 |
| 1 | 31 | ENSG00000130635 | COL5A1 | 2.23 |
| 1 | 32 | ENSG00000257017 | HP | 2.2 |
| 1 | 33 | ENSG00000115221 | ITGB6 | 2.2 |
| 1 | 34 | ENSG00000204262 | COL5A2 | 2.19 |
| 1 | 35 | ENSG00000137312 | FLOT1 | 2.18 |
| 1 | 36 | ENSG00000140254 | DUOXA1 | 2.17 |
| 1 | 37 | ENSG00000096060 | FKBP5 | 2.16 |
| 1 | 38 | ENSG00000189377 | CXCL17 | 2.13 |
| 1 | 39 | ENSG00000123500 | COL10A1 | 2.1 |
| 1 | 40 | ENSG00000272398 | CD24 | 2.06 |
| 1 | 41 | ENSG00000143341 | HMCN1 | 2.06 |
| 1 | 42 | ENSG00000187955 | COL14A1 | 2.06 |

| | | | | |
|---|---|---|---|---|
| 1 | 43 | ENSG00000136352 | NKX2-1 | 2.04 |
| 1 | 44 | ENSG00000062038 | CDH3 | 2.03 |
| 1 | 45 | ENSG00000158467 | AHCYL2 | 1.99 |
| 1 | 46 | ENSG00000127249 | ATP13A4 | 1.99 |
| 1 | 47 | ENSG00000184012 | TMPRSS2 | 1.98 |
| 1 | 48 | ENSG00000128422 | KRT17 | 1.98 |
| 1 | 49 | ENSG00000100033 | PRODH | 1.97 |
| 1 | 50 | ENSG00000162896 | PIGR | 1.97 |
| 2 | 1 | ENSG00000168484 | SFTPC | 4.77 |
| 2 | 2 | ENSG00000122852 | SFTPA1 | 4 |
| 2 | 3 | ENSG00000185303 | SFTPA2 | 3.96 |
| 2 | 4 | ENSG00000096088 | PGC | 3.48 |
| 2 | 5 | ENSG00000131400 | NAPSA | 3.33 |
| 2 | 6 | ENSG00000124107 | SLPI | 3.23 |
| 2 | 7 | ENSG00000133661 | SFTPD | 3.1 |
| 2 | 8 | ENSG00000197249 | SERPINA1 | 3.09 |
| 2 | 9 | ENSG00000205364 | MT1M | 3.07 |
| 2 | 10 | ENSG00000135480 | KRT7 | 3 |
| 2 | 11 | ENSG00000171476 | HOPX | 2.89 |
| 2 | 12 | ENSG00000188536 | HBA2 | 2.84 |
| 2 | 13 | ENSG00000170421 | KRT8 | 2.81 |
| 2 | 14 | ENSG00000163682 | RPL9 | 2.78 |
| 2 | 15 | ENSG00000111057 | KRT18 | 2.77 |
| 2 | 16 | ENSG00000171345 | KRT19 | 2.74 |
| 2 | 17 | ENSG00000175793 | SFN | 2.73 |
| 2 | 18 | ENSG00000130208 | APOC1 | 2.72 |
| 2 | 19 | ENSG00000171557 | FGG | 2.68 |
| 2 | 20 | ENSG00000090339 | ICAM1 | 2.66 |
| 2 | 21 | ENSG00000090382 | LYZ | 2.63 |
| 2 | 22 | ENSG00000066405 | CLDN18 | 2.61 |
| 2 | 23 | ENSG00000134020 | PEBP4 | 2.57 |
| 2 | 24 | ENSG00000198502 | HLA-DRB5 | 2.55 |
| 2 | 25 | ENSG00000013588 | GPRC5A | 2.55 |
| 2 | 26 | ENSG00000170323 | FABP4 | 2.48 |
| 2 | 27 | ENSG00000110195 | FOLR1 | 2.45 |
| 2 | 28 | ENSG00000167972 | ABCA3 | 2.44 |
| 2 | 29 | ENSG00000170786 | SDR16C5 | 2.44 |
| 2 | 30 | ENSG00000138821 | SLC39A8 | 2.42 |
| 2 | 31 | ENSG00000148671 | ADIRF | 2.4 |
| 2 | 32 | ENSG00000114854 | TNNC1 | 2.38 |
| 2 | 33 | ENSG00000148677 | ANKRD1 | 2.37 |
| 2 | 34 | ENSG00000162551 | ALPL | 2.35 |
| 2 | 35 | ENSG00000078081 | LAMP3 | 2.34 |
| 2 | 36 | ENSG00000163435 | ELF3 | 2.32 |
| 2 | 37 | ENSG00000172602 | RND1 | 2.32 |
| 2 | 38 | ENSG00000184292 | TACSTD2 | 2.32 |
| 2 | 39 | ENSG00000142973 | CYP4B1 | 2.32 |
| 2 | 40 | ENSG00000099994 | SUSD2 | 2.32 |

| | | | | |
|---|---|---|---|---|
| 2 | 41 | ENSG00000168309 | FAM107A | 2.31 |
| 2 | 42 | ENSG00000189143 | CLDN4 | 2.3 |
| 2 | 43 | ENSG00000268104 | SLC6A14 | 2.28 |
| 2 | 44 | ENSG00000138772 | ANXA3 | 2.27 |
| 2 | 45 | ENSG00000165272 | AQP3 | 2.25 |
| 2 | 46 | ENSG00000086548 | CEACAM6 | 2.25 |
| 2 | 47 | ENSG00000169583 | CLIC3 | 2.24 |
| 2 | 48 | ENSG00000086544 | ITPKC | 2.24 |
| 2 | 49 | ENSG00000095713 | CRTAC1 | 2.21 |
| 2 | 50 | ENSG00000183607 | GKN2 | 2.2 |
| 3 | 1 | ENSG00000211651 | IGLV1-44 | 6.89 |
| 3 | 2 | ENSG00000211592 | IGKC | 6.72 |
| 3 | 3 | ENSG00000211677 | IGLC2 | 6.58 |
| 3 | 4 | ENSG00000211893 | IGHG2 | 6.42 |
| 3 | 5 | ENSG00000211679 | IGLC3 | 6.41 |
| 3 | 6 | ENSG00000211897 | IGHG3 | 6.32 |
| 3 | 7 | ENSG00000251546 | IGKV1D-39 | 6.22 |
| 3 | 8 | ENSG00000211666 | IGLV2-14 | 6.09 |
| 3 | 9 | ENSG00000211675 | IGLC1 | 6.08 |
| 3 | 10 | ENSG00000211648 | IGLV1-47 | 5.91 |
| 3 | 11 | ENSG00000211598 | IGKV4-1 | 5.88 |
| 3 | 12 | ENSG00000278196 | IGLV2-8 | 5.84 |
| 3 | 13 | ENSG00000211668 | IGLV2-11 | 5.79 |
| 3 | 14 | ENSG00000211660 | IGLV2-23 | 5.74 |
| 3 | 15 | ENSG00000211663 | IGLV3-19 | 5.73 |
| 3 | 16 | ENSG00000211659 | IGLV3-25 | 5.7 |
| 3 | 17 | ENSG00000211673 | IGLV3-1 | 5.64 |
| 3 | 18 | ENSG00000211669 | IGLV3-10 | 5.54 |
| 3 | 19 | ENSG00000254709 | IGLL5 | 5.49 |
| 3 | 20 | ENSG00000211955 | IGHV3-33 | 5.47 |
| 3 | 21 | ENSG00000211662 | IGLV3-21 | 5.44 |
| 3 | 22 | ENSG00000275063 | AC233755.1 | 5.37 |
| 3 | 23 | ENSG00000211640 | IGLV6-57 | 5.34 |
| 3 | 24 | ENSG00000132465 | JCHAIN | 5.32 |
| 3 | 25 | ENSG00000211644 | IGLV1-51 | 5.32 |
| 3 | 26 | ENSG00000224041 | IGKV3D-15 | 5.19 |
| 3 | 27 | ENSG00000231475 | IGHV4-31 | 5.12 |
| 3 | 28 | ENSG00000211685 | IGLC7 | 4.93 |
| 3 | 29 | ENSG00000242534 | IGKV2D-28 | 4.9 |
| 3 | 30 | ENSG00000211649 | IGLV7-46 | 4.87 |
| 3 | 31 | ENSG00000211638 | IGLV8-61 | 4.84 |
| 3 | 32 | ENSG00000211670 | IGLV3-9 | 4.78 |
| 3 | 33 | ENSG00000211637 | IGLV4-69 | 4.74 |
| 3 | 34 | ENSG00000274576 | IGHV2-70 | 4.72 |
| 3 | 35 | ENSG00000243264 | IGKV2D-29 | 4.5 |
| 3 | 36 | ENSG00000211625 | IGKV3D-20 | 4.46 |
| 3 | 37 | ENSG00000277856 | AC233755.2 | 4.37 |
| 3 | 38 | ENSG00000170476 | MZB1 | 4.33 |

| | | | | |
|---|---|---|---|---|
| 3 | 39 | ENSG00000211658 | IGLV3-27 | 4.2 |
| 3 | 40 | ENSG00000276566 | IGKV1D-13 | 4.05 |
| 3 | 41 | ENSG00000280411 | IGHV1-69D | 4.05 |
| 3 | 42 | ENSG00000211664 | IGLV2-18 | 3.98 |
| 3 | 43 | ENSG00000211632 | IGKV3D-11 | 3.96 |
| 3 | 44 | ENSG00000232216 | IGHV3-43 | 3.88 |
| 3 | 45 | ENSG00000277836 | AC141272.1 | 3.82 |
| 3 | 46 | ENSG00000211642 | IGLV10-54 | 3.79 |
| 3 | 47 | ENSG00000233999 | IGKV3OR2-268 | 3.66 |
| 3 | 48 | ENSG00000211898 | IGHD | 3.65 |
| 3 | 49 | ENSG00000251039 | IGKV2D-40 | 3.65 |
| 3 | 50 | ENSG00000241244 | IGKV1D-16 | 3.64 |
| 4 | 1 | ENSG00000206503 | HLA-A | 4.38 |
| 4 | 2 | ENSG00000129824 | RPS4Y1 | 3.83 |
| 4 | 3 | ENSG00000166292 | TMEM100 | 3.8 |
| 4 | 4 | ENSG00000108576 | SLC6A4 | 3.71 |
| 4 | 5 | ENSG00000204525 | HLA-C | 3.69 |
| 4 | 6 | ENSG00000142748 | FCN3 | 3.57 |
| 4 | 7 | ENSG00000234745 | HLA-B | 3.51 |
| 4 | 8 | ENSG00000240247 | DEFA1B | 3.28 |
| 4 | 9 | ENSG00000204642 | HLA-F | 3.28 |
| 4 | 10 | ENSG00000204592 | HLA-E | 3.21 |
| 4 | 11 | ENSG00000143546 | S100A8 | 3.1 |
| 4 | 12 | ENSG00000116016 | EPAS1 | 2.97 |
| 4 | 13 | ENSG00000114812 | VIPR1 | 2.95 |
| 4 | 14 | ENSG00000122679 | RAMP3 | 2.94 |
| 4 | 15 | ENSG00000105374 | NKG7 | 2.92 |
| 4 | 16 | ENSG00000163220 | S100A9 | 2.91 |
| 4 | 17 | ENSG00000204389 | HSPA1A | 2.91 |
| 4 | 18 | ENSG00000131477 | RAMP2 | 2.87 |
| 4 | 19 | ENSG00000163815 | CLEC3B | 2.87 |
| 4 | 20 | ENSG00000165810 | BTNL9 | 2.84 |
| 4 | 21 | ENSG00000244734 | HBB | 2.8 |
| 4 | 22 | ENSG00000184113 | CLDN5 | 2.79 |
| 4 | 23 | ENSG00000121858 | TNFSF10 | 2.77 |
| 4 | 24 | ENSG00000102760 | RGCC | 2.76 |
| 4 | 25 | ENSG00000167434 | CA4 | 2.76 |
| 4 | 26 | ENSG00000137834 | SMAD6 | 2.75 |
| 4 | 27 | ENSG00000206172 | HBA1 | 2.72 |
| 4 | 28 | ENSG00000103034 | NDRG4 | 2.7 |
| 4 | 29 | ENSG00000131203 | IDO1 | 2.69 |
| 4 | 30 | ENSG00000067048 | DDX3Y | 2.68 |
| 4 | 31 | ENSG00000271503 | CCL5 | 2.64 |
| 4 | 32 | ENSG00000139567 | ACVRL1 | 2.63 |
| 4 | 33 | ENSG00000162654 | GBP4 | 2.63 |
| 4 | 34 | ENSG00000138755 | CXCL9 | 2.6 |
| 4 | 35 | ENSG00000163221 | S100A12 | 2.59 |
| 4 | 36 | ENSG00000105227 | PRX | 2.58 |

| | | | | |
|---|---|---|---|---|
| 4 | 37 | ENSG00000196260 | SFTA2 | 2.58 |
| 4 | 38 | ENSG00000064989 | CALCRL | 2.57 |
| 4 | 39 | ENSG00000010319 | SEMA3G | 2.57 |
| 4 | 40 | ENSG00000263155 | MYZAP | 2.55 |
| 4 | 41 | ENSG00000115523 | GNLY | 2.54 |
| 4 | 42 | ENSG00000204305 | AGER | 2.52 |
| 4 | 43 | ENSG00000198692 | EIF1AY | 2.52 |
| 4 | 44 | ENSG00000100450 | GZMH | 2.49 |
| 4 | 45 | ENSG00000164120 | HPGD | 2.46 |
| 4 | 46 | ENSG00000168685 | IL7R | 2.45 |
| 4 | 47 | ENSG00000239839 | DEFA3 | 2.43 |
| 4 | 48 | ENSG00000169245 | CXCL10 | 2.43 |
| 4 | 49 | ENSG00000242574 | HLA-DMB | 2.41 |
| 4 | 50 | ENSG00000012817 | KDM5D | 2.41 |
| 5 | 1 | ENSG00000125999 | BPIFB1 | 4.86 |
| 5 | 2 | ENSG00000149021 | SCGB1A1 | 4.83 |
| 5 | 3 | ENSG00000124237 | C20orf85 | 3.6 |
| 5 | 4 | ENSG00000164972 | C9orf24 | 3.56 |
| 5 | 5 | ENSG00000160472 | TMEM190 | 3.37 |
| 5 | 6 | ENSG00000166959 | MS4A8 | 3.35 |
| 5 | 7 | ENSG00000186081 | KRT5 | 3.28 |
| 5 | 8 | ENSG00000179902 | C1orf194 | 3.22 |
| 5 | 9 | ENSG00000186973 | FAM183A | 3.11 |
| 5 | 10 | ENSG00000105519 | CAPS | 3.11 |
| 5 | 11 | ENSG00000160188 | RSPH1 | 3.03 |
| 5 | 12 | ENSG00000188817 | SNTN | 2.99 |
| 5 | 13 | ENSG00000148735 | PLEKHS1 | 2.93 |
| 5 | 14 | ENSG00000185681 | MORN5 | 2.93 |
| 5 | 15 | ENSG00000183644 | HOATZ | 2.91 |
| 5 | 16 | ENSG00000161055 | SCGB3A1 | 2.91 |
| 5 | 17 | ENSG00000117472 | TSPAN1 | 2.88 |
| 5 | 18 | ENSG00000152611 | CAPSL | 2.86 |
| 5 | 19 | ENSG00000161905 | ALOX15 | 2.82 |
| 5 | 20 | ENSG00000034239 | EFCAB1 | 2.82 |
| 5 | 21 | ENSG00000153789 | CIBAR2 | 2.81 |
| 5 | 22 | ENSG00000263639 | MSMB | 2.8 |
| 5 | 23 | ENSG00000136918 | WDR38 | 2.76 |
| 5 | 24 | ENSG00000243955 | GSTA1 | 2.74 |
| 5 | 25 | ENSG00000188931 | CFAP126 | 2.73 |
| 5 | 26 | ENSG00000231738 | TSPAN19 | 2.68 |
| 5 | 27 | ENSG00000186710 | CFAP73 | 2.68 |
| 5 | 28 | ENSG00000277639 | CHD9NB | 2.67 |
| 5 | 29 | ENSG00000198183 | BPIFA1 | 2.67 |
| 5 | 30 | ENSG00000004838 | ZMYND10 | 2.66 |
| 5 | 31 | ENSG00000162004 | CCDC78 | 2.63 |
| 5 | 32 | ENSG00000167858 | TEKT1 | 2.62 |
| 5 | 33 | ENSG00000166596 | CFAP52 | 2.61 |
| 5 | 34 | ENSG00000179813 | FAM216B | 2.58 |

| | | | | |
|---|---|---|---|---|
| 5 | 35 | ENSG00000145491 | ROPN1L | 2.57 |
| 5 | 36 | ENSG00000077327 | SPAG6 | 2.56 |
| 5 | 37 | ENSG00000215187 | FAM166B | 2.53 |
| 5 | 38 | ENSG00000133665 | DYDC2 | 2.5 |
| 5 | 39 | ENSG00000203985 | LDLRAD1 | 2.49 |
| 5 | 40 | ENSG00000129654 | FOXJ1 | 2.48 |
| 5 | 41 | ENSG00000204711 | C9orf135 | 2.44 |
| 5 | 42 | ENSG00000165309 | ARMC3 | 2.44 |
| 5 | 43 | ENSG00000167653 | PSCA | 2.43 |
| 5 | 44 | ENSG00000174898 | CATSPERD | 2.43 |
| 5 | 45 | ENSG00000188659 | SAXO2 | 2.42 |
| 5 | 46 | ENSG00000148346 | LCN2 | 2.41 |
| 5 | 47 | ENSG00000122735 | DNAI1 | 2.41 |
| 5 | 48 | ENSG00000141294 | LRRC46 | 2.41 |
| 5 | 49 | ENSG00000197748 | CFAP43 | 2.38 |
| 5 | 50 | ENSG00000112539 | C6orf118 | 2.38 |
| 6 | 1 | ENSG00000136244 | IL6 | 4.53 |
| 6 | 2 | ENSG00000158859 | ADAMTS4 | 4.21 |
| 6 | 3 | ENSG00000108691 | CCL2 | 4.16 |
| 6 | 4 | ENSG00000163661 | PTX3 | 4.11 |
| 6 | 5 | ENSG00000125148 | MT2A | 4.05 |
| 6 | 6 | ENSG00000169429 | CXCL8 | 3.81 |
| 6 | 7 | ENSG00000106366 | SERPINE1 | 3.7 |
| 6 | 8 | ENSG00000175592 | FOSL1 | 3.66 |
| 6 | 9 | ENSG00000112096 | SOD2 | 3.64 |
| 6 | 10 | ENSG00000102265 | TIMP1 | 3.61 |
| 6 | 11 | ENSG00000134531 | EMP1 | 3.59 |
| 6 | 12 | ENSG00000205362 | MT1A | 3.56 |
| 6 | 13 | ENSG00000102802 | MEDAG | 3.54 |
| 6 | 14 | ENSG00000187193 | MT1X | 3.53 |
| 6 | 15 | ENSG00000108342 | CSF3 | 3.49 |
| 6 | 16 | ENSG00000128342 | LIF | 3.42 |
| 6 | 17 | ENSG00000137801 | THBS1 | 3.32 |
| 6 | 18 | ENSG00000184557 | SOCS3 | 3.27 |
| 6 | 19 | ENSG00000196136 | SERPINA3 | 3.23 |
| 6 | 20 | ENSG00000123689 | G0S2 | 3.21 |
| 6 | 21 | ENSG00000167772 | ANGPTL4 | 3.16 |
| 6 | 22 | ENSG00000148926 | ADM | 3.09 |
| 6 | 23 | ENSG00000132510 | KDM6B | 3.07 |
| 6 | 24 | ENSG00000136997 | MYC | 3.06 |
| 6 | 25 | ENSG00000131459 | GFPT2 | 3.05 |
| 6 | 26 | ENSG00000125144 | MT1G | 2.99 |
| 6 | 27 | ENSG00000104635 | SLC39A14 | 2.97 |
| 6 | 28 | ENSG00000169715 | MT1E | 2.96 |
| 6 | 29 | ENSG00000187479 | C11orf96 | 2.96 |
| 6 | 30 | ENSG00000188257 | PLA2G2A | 2.94 |
| 6 | 31 | ENSG00000163638 | ADAMTS9 | 2.93 |
| 6 | 32 | ENSG00000081041 | CXCL2 | 2.93 |

| | | | | |
|---|---|---|---|---|
| 6 | 33 | ENSG00000115009 | CCL20 | 2.91 |
| 6 | 34 | ENSG00000103196 | CRISPLD2 | 2.91 |
| 6 | 35 | ENSG00000128016 | ZFP36 | 2.89 |
| 6 | 36 | ENSG00000213088 | ACKR1 | 2.87 |
| 6 | 37 | ENSG00000133800 | LYVE1 | 2.83 |
| 6 | 38 | ENSG00000142871 | CCN1 | 2.83 |
| 6 | 39 | ENSG00000123342 | MMP19 | 2.82 |
| 6 | 40 | ENSG00000125740 | FOSB | 2.81 |
| 6 | 41 | ENSG00000175084 | DES | 2.76 |
| 6 | 42 | ENSG00000243509 | TNFRSF6B | 2.73 |
| 6 | 43 | ENSG00000166741 | NNMT | 2.72 |
| 6 | 44 | ENSG00000006327 | TNFRSF12A | 2.71 |
| 6 | 45 | ENSG00000163739 | CXCL1 | 2.71 |
| 6 | 46 | ENSG00000100292 | HMOX1 | 2.68 |
| 6 | 47 | ENSG00000099860 | GADD45B | 2.66 |
| 6 | 48 | ENSG00000185022 | MAFF | 2.62 |
| 6 | 49 | ENSG00000104368 | PLAT | 2.56 |
| 6 | 50 | ENSG00000105825 | TFPI2 | 2.53 |
